# Secreted protein methyltransferase METTL9 catalyzes *N*π-histidine methylation of extracellular plasma proteins

**DOI:** 10.64898/2025.12.23.696137

**Authors:** Neo Ikeda, Hiroaki Daitoku, Natsuki Odate, Naoki Sekiguchi, Ryuhei Tsukamoto, Rikuto Kondo, Song-iee Han, Kaori Motomura, Koichiro Kako, Akiyoshi Fukamizu

## Abstract

Post-translational modifications (PTMs) of proteins alter their biophysical properties, thus affecting their activity, localization and interactions. These reactions are principally intracellular events; however, to date, only phosphorylation has been shown to occur within the extracellular space. Here, we identified METTL9 as the first secreted methyltransferase responsible for *N*π-histidine methylation. METTL9 undergoes N-linked glycosylation, thereby forming dimers via disulfide bonds. Using a split-luciferase complementary assay, we revealed that N-glycosylated METTL9 is secreted extracellularly via the ER-Golgi pathway. Endogenous METTL9 is highly expressed in HL60 cells during neutrophil-like differentiation and secreted extracellularly. METTL9 catalyzes *N*π-methylhistidine formation in plasma proteins, in which the thyroxine transporter transthyretin (TTR) and copper transporter ceruloplasmin are identified as substrates for methylation *in vitro*. Both methylations occur at the His-x-His motif, a recognition sequence for METTL9, and TTR methylation decreases its binding affinity to zinc. Our results establish that histidine methylation is the second extracellular PTM, following phosphorylation.

## Introduction

Post-translational modifications (PTMs) of proteins regulate almost all aspects of cellular processes by rapidly altering the functional properties of proteins, including changes in their enzymatic activity, subcellular localization, interaction partners, protein stability, DNA binding, and interaction partners [1]. The list of known PTMs is quite extensive, but common PTMs can be broadly classified into three categories depending on the molecule attached to the target proteins: first, the addition of chemical groups, such as phosphorylation [2], acetylation [3], and methylation [4]; second, conjugation with small proteins, as in ubiquitination and SUMOylation [5] [6]; and third, the attachment of sugar chains (glycans), as in *N*- and *O*-linked glycosylation [7]. Despite being one of the smallest modifications, methylation (the addition of -CH_3_ groups from S-adenosyl-L-methionine) is probably the most widespread and functionally diverse PTMs. Protein methylation commonly occurs on the side chain of lysine and arginine residues, producing three distinct methylated forms: mono-, di-, and tri-methylation of lysine and mono-, symmetric di-, and asymmetric di-methylation of arginine. Lysine methylation exists particularly in histones and plays essential roles in regulating chromatin dynamics [8], whereas arginine methylation is found in diverse proteins and is involved in a wide range of cellular events, including transcription, RNA processing, signal transduction, and DNA damage response [9]. In addition to lysine and arginine, the methylation of another basic amino acid, histidine, has recently attracted attention [10].

Histidine residue can potentially be monomethylated on the nitrogen in either position 1 (*N*π) or 3 (*N*τ) of the imidazole ring, yielding *N*π-methylhistidine (also referred to as 1MH) or *N*τ-methylhistidine (3MH), respectively. Although protein histidine methylation was first found as 3MH in the constituents of actin and myosin from muscle proteins more than half a century ago [11, 12], the responsible enzyme has long remained unknown. However, starting with the first identification of the histidine methyltransferase (MTase) Hpm-1 in yeast [13], a series of mammalian MTases have recently been discovered, namely SETD3 [14, 15], METTL18 [16], METTL9 [17–19], and CARNMT1 [20]. Among these, METTL9 is the MTase that introduces 1MH at the His-x-His (where x denotes a small side-chain residue) motif-containing proteins, such as the proinflammatory protein S100A9 and the NDUFB3 subunit of mitochondrial respiratory Complex I [17–19, 21]. Importantly, we and another group have revealed that the presence of 1MH in an HxH-containing peptide decreases its binding affinity to Zn^2+^, indicating a functional consequence of protein histidine methylation [17, 18]. In addition, we previously found that METTL9 localizes to the endoplasmic reticulum when overexpressed in mammalian cultured cells; however, its biological significance remains unknown [18].

In most cases, the PTMs described above are intracellular reactions, and their modifying enzymes are co-localized with and then encounter substrate proteins in the cytoplasm and organelles; however, only “phosphorylation” is known to occur in the extracellular space [22, 23]. The first kinase that phosphorylates protein domains destined to be extracellular is the four-jointed (Fj) in *Drosophila* [24]. Fj is a Golgi-residing, single-pass transmembrane protein that phosphorylates serine or threonine residues within the extracellular cadherin domains of Fat and its transmembrane ligand, Dachsous [24]. In contrast, in mammals, Fam20C, a family of atypical protein kinases localized in the Golgi, has been identified as the “real kinase” for casein, which is abundant in milk and the first phosphoprotein reported over 140 years ago [25]. Fam20C is also a secreted protein that phosphorylates S-x-E/pS motifs within secretory pathway proteins involved in biomineralization [25]. Additionally, vertebrate lonesome kinase (VLK) has been shown to be a secreted protein responsible for extracellular tyrosine phosphorylation [26]. VLK is highly expressed in platelets and can be rapidly and quantitatively secreted from platelets in response to various stimuli [26]. Thus far, the discovery of secreted kinases has gradually shed light on the existence of extracellular protein phosphorylation; however, there are no reports of secreted forms of modifying enzymes other than kinases. Here, we identified METTL9 as the first secreted methyltransferase that methylates the plasma proteins transthyretin and ceruloplasmin. Our findings suggest that extracellular PTMs represent a universal mechanism that extends beyond the scope of phosphorylation.

## Results

### N-linked glycosylation occurs in mouse METTL9 at Asn-35 and Asn-86

METTL9 proteins across species contain a putative signal peptide (SP) sequence composed of abundant hydrophobic amino acids in the N-terminal region (Figure 1A). Thus, we predicted the presence of SPs and cleavage sites of METTL9 orthologs using the SignalP server [27]. Consistent with our previous report [18], SPs are conserved in mammalian METTL9, including in *Homo sapiens* and *Mus musculus*, but not in other orthologs, such as *Danio rerio*, *Drosophila melanogaster*, and *Caenorhabditis elegans* (Figure S1A-S1E). Meanwhile, in the case of mouse METTL9, Western blotting showed two extra bands above the calculated molecular weight of 35 kDa when expressed as C-terminal FLAG-tagged proteins in HEK293T cells, whereas deletion of the SP sequence from the 1st to 18th amino acids abolished the additional bands (Figure 1B). The bottom band below the 35 kDa marker is assumed to be a product that was cleaved longer than the predicted SP sequence (* in Figure 1B).

**Figure 1.**
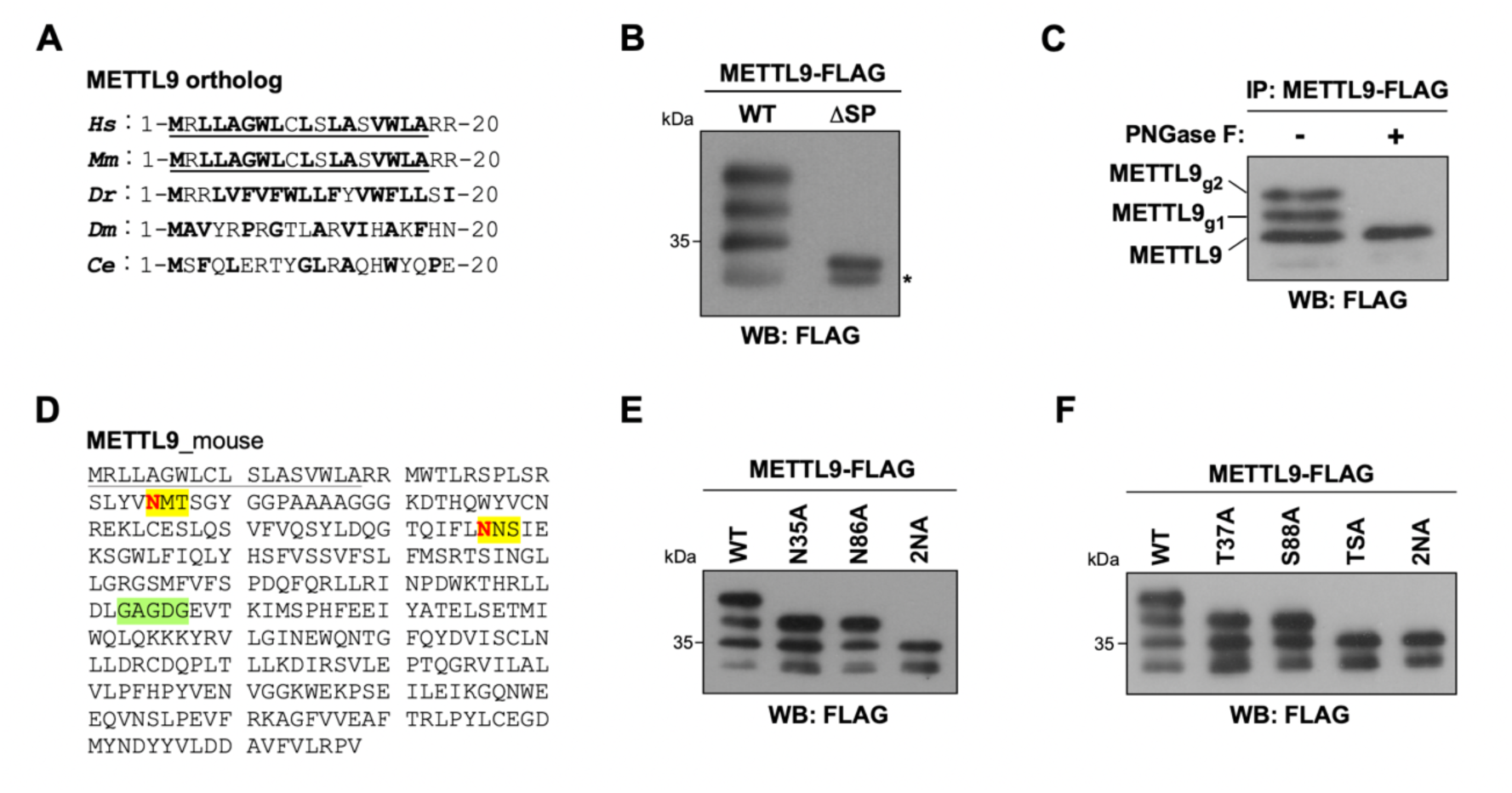
METTL9 is N-glycosylated at Asn-35 and Asn-86. **(A)** Sequence alignment of the N-terminal end of METTL9 orthologs across species. *Hs*, *Homo sapiens*; *Mm*, *Mus musculus*; *Dr*, *Danio rerio*; *Dm*, *Drosophila melanogaster*; *Ce*, *Caenorhabditis elegans*. The SignalP-predicted signal peptide (SP) sequences are underlined. Hydrophobic amino acids are shown in bold. (**B**) Band patterns of METTL9 WT and ΔSP mutant. Cell lysates expressing METTL9-FLAG WT or ΔSP mutant were analyzed by Western blotting with an anti-FLAG antibody. An asterisk indicates the band corresponding to the further cleaved METTL9. (**C**) Digestion of METTL9 with a deglycosylation enzyme. Immunoprecipitated METTL9-FLAG was incubated with (+) or without (-) PNGase F to cleave N-glycans and then analyzed by Western blotting with an anti-FLAG antibody. (**D**) Amino acid sequence of mouse METTL9. The consensus motif for N-glycosylation (N-X-S/T) and the SAM-binding motif (GxGxG) are highlighted in yellow and green, respectively. Asparagine residues at positions 35 and 86 are denoted in bold red. Predicted SP sequences are underlined. (**E** and **F**) Band patterns of METTL9 WT and N-glycosylation-deficient mutants. Cell lysates expressing METTL9-FLAG WT or N-X-S/T sequon mutants were analyzed by Western blotting with an anti-FLAG antibody.

Considering the endoplasmic reticulum (ER) localization of METTL9 [18], we hypothesized that METTL9 undergoes N-linked glycosylation in an SP-dependent manner. To test this hypothesis, we used PNGase F, which cleaves the carboxamido group of N-glycosylated asparagine residues. As expected, the two shift-up bands of immunoprecipitated METTL9 were eliminated by this digestion (Figure 1C), suggesting that N-glycans are added to two distinct asparagine residues of METTL9 in cells. Supporting this result, the consensus sequence of N-linked glycosylation (N-X-S/T sequon, where X denotes any amino acid except proline) exists in two locations in mouse METTL9 (Figure 1D). Furthermore, when the asparagine residues within the two motifs (Asn-35 and Asn-86) were replaced with alanine, the top and second bands disappeared in the single mutants (N35A or N86A) and the double mutant (2NA), respectively (Figure 1E). Similar results were obtained by substituting Thr-37 or Ser-88 in the sequon with alanine (Figure 1F). These data indicate that METTL9 is N-glycosylated at Asn-35 and Asn-86 in an SP-dependent manner.

### N-glycosylation of METTL9 is required for dimerization via disulfide bond formation

To investigate the functional significance of N-glycosylation for METTL9, we first compared the methyltransferase activities of wild-type and 2NA mutant METTL9 expressed in HEK293T cells using recombinant S100A9 as a substrate. However, unlike the 2GA mutant, which is defective in SAM binding, the 2NA mutant methylated S100A9 to the same extent as the wild-type protein *in vitro* (Figure 2A). Next, to determine whether N-glycosylation of METTL9 is involved in protein-protein interactions, the co-immunoprecipitated proteins with METTL9-FLAG were analyzed by MALDI-TOF/MS. We identified calnexin, a lectin chaperone essential for protein folding in the ER, as a binding partner of METTL9 wild type but not the 2NA mutant (Figure 2B and S2A). Although calnexin has been reported to interact with GFP-fused METTL9 [28], we further revealed that N-glycosylation is required for the interaction between METTL9 and calnexin (Figure 2C). In contrast, N-glycosylation was not essential for the ER localization of METTL9 (Figure S2B). In addition, because calnexin and its soluble homolog, calreticulin, often serve as a scaffold bridging together N-glycosylated proteins with other ER chaperones, such as ERp57, we tested whether ERp57 was also included in the immunoprecipitates of METTL9. Consistent with calnexin, METTL9 showed glycan-dependent binding to ERp57 (Figure 2D).

**Figure 2.**
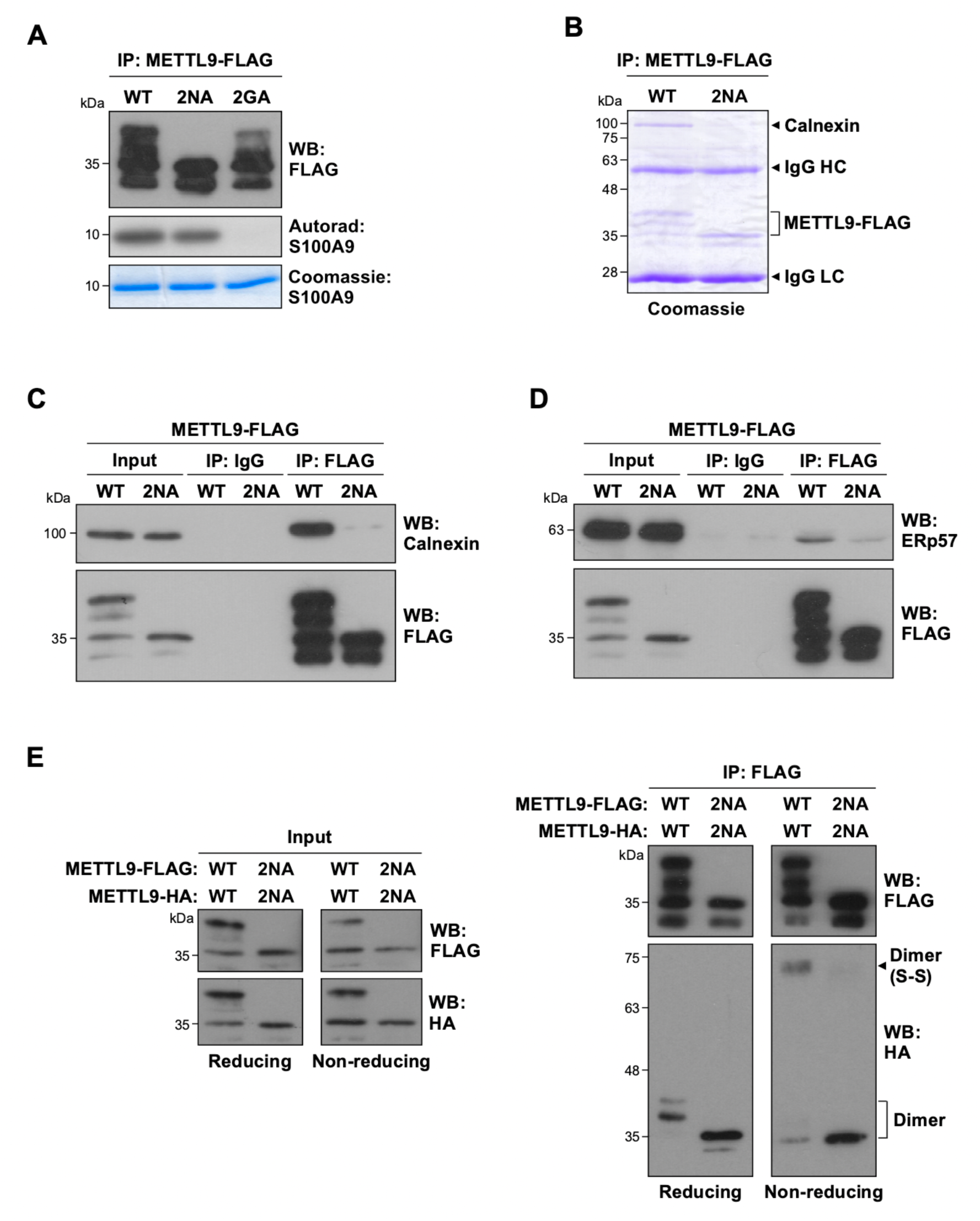
METTL9 forms a disulfide bond-mediated dimer depending on N-glycosylation. **(A)** N-glycosylation of METTL9 has no effect on its methyltransferase activity. Immunoprecipitated METTL9-FLAG WT, 2NA, and 2GA mutants were incubated with recombinant S100A9 and ^3^H-labeled SAM. METTL9-FLAG and S100A9 in the reaction were detected by Western blotting (*Top*) and Coomassie blue staining (*Bottom*), respectively. The incorporation of [^3^H]-methyl into S100A9 was visualized by autoradiography (*Middle*). (**B**) Binding proteins of N-glycosylated METTL9. Co-immunoprecipitated proteins with METTL9-FLAG WT or 2NA mutant in HEK293T cells were stained with Coomassie blue. A band around 100 kDa was identified as Calnexin by MALDI-TOF/MS analysis. (**C** and **D**) Calnexin and ERp57 bind to N-glycosylated METTL9. Immunoprecipitated METTL9-FLAG WT or 2NA mutant proteins were analyzed by Western blotting with anti-calnexin (**C**), anti-ERp57 (**D**), and anti-FLAG antibodies. Normal mouse IgG was used as the negative control. The input indicates 2% of the whole-cell lysates used for immunoprecipitation. (**E**) Dimerization of N-glycosylated METTL9 via disulfide bond formation in cells. Cell lysates co-expressing FLAG- and HA-tagged METTL9, as indicated, were immunoprecipitated with an anti-FLAG antibody under reducing or non-reducing conditions. Input indicates 2% of the whole-cell lysates used for immunoprecipitation (*Left*). Immunoprecipitated METTL9-FLAG WT or 2NA mutant was analyzed by Western blotting with anti-FLAG and anti-HA antibodies (*Right*). The arrowhead indicates a band of METTL9 dimers with disulfide bond formation. The bracket indicates the bands of the METTL9 dimer without disulfide bond formation.

Given the following two points: first, methyltransferases often form dimers to retain SAM in the active site [29], properly recognize substrates [30], or maintain their own stability [31]; and second, ERp57 is a glycoprotein-specific thiol-disulfide oxidoreductase [=protein disulfide isomerase] [32], it is possible that METTL9 might dimerize via disulfide bond formation with the help of ERp57 in the ER. To test this possibility, we performed a co-immunoprecipitation assay between FLAG- and HA-tagged METTL9, followed by SDS-PAGE under reducing and non-reducing conditions. Western blotting of reducing gel showed bands of dimers at an apparent molecular weight of ∼35 kDa, indicating that both glycosylated and non-glycosylated METTL9 form dimers (Figure 2E, right). Remarkably, a band was detected at a height considered to be a dimer (∼70 kDa) under non-reducing conditions (Figure 2E, right). Overall, these data suggest that N-glycosylated METTL9 can dimerize through disulfide bonds in the ER.

### N-glycosylated METTL9 is secreted to the extracellular space

The finding that *N*-glycans are added to METTL9 prompted us to examine the possibility of its extracellular secretion, as well as that of most N-glycosylated proteins. To this end, we employed the Nano-Glo® HiBiT Extracellular Detection System, in which the split version of a small nanoluciferase fragment (HiBiT) is tagged to a protein of interest and expressed in cells; thus, its extracellular secretion can be measured in the culture medium by nanoluciferase complementation (Figure 3A) [33]. We expressed C-terminal HiBiT-tagged METTL9 proteins in HEK293T cells and monitored their amounts outside the cells as changes in luminescence of the culture medium. Luminescence levels gradually increased over time after medium change; however, treatment with brefeldin A, an inhibitor of protein transport from the ER to the Golgi membrane, completely suppressed the increase in luminescent signals (Figure 3B). Importantly, the ratio of extracellular to lytic signals at post-6 hours substantially decreased (Figure 3C), indicating that METTL9-HiBiT is secreted through the classical ER-Golgi pathway. We further examined the role of N-glycosylation in METTL9 secretion using HiBiT-tagged 2NA and ΔSP mutant proteins. In contrast to wild-type METTL9, both 2NA and ΔSP mutants failed to produce extracellular luminescent signals (Figure 3D and E), despite similar expression levels of the two mutants (Figure 3F).

**Figure 3.**
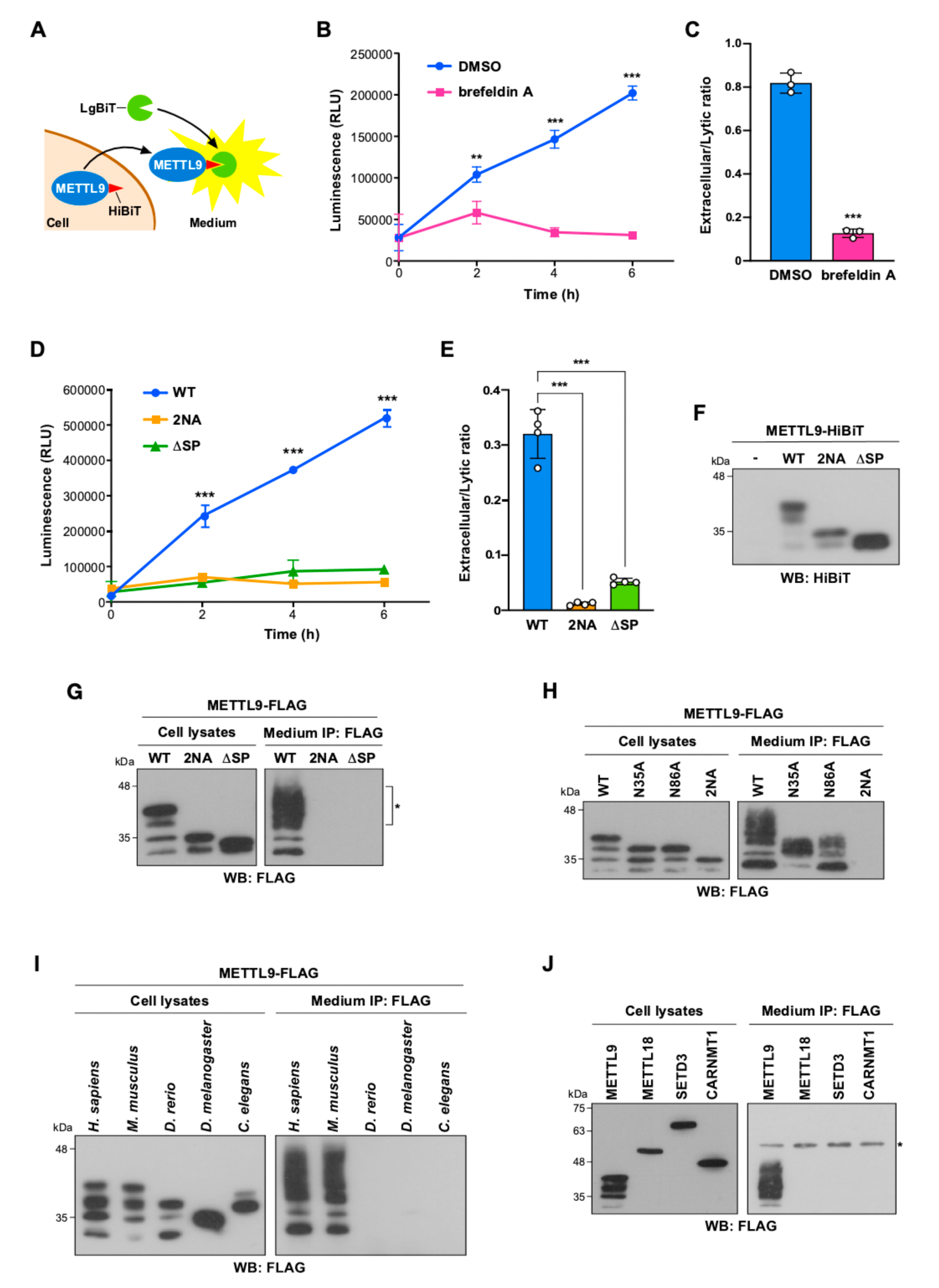
N-glycosylated METTL9 is secreted extracellularly. **(A)** Schematic representation of the HiBiT extracellular detection system. HiBiT-tagged METTL9 is complemented with LgBiT in the culture medium and then detected as a fluorescent signal. (**B** and **C**) Inhibitory effect of brefeldin A on METTL9 secretion. Luciferase activities of METTL9-HiBiT treated with DMSO or brefeldin A were plotted as line graphs over time (**B**). The ratio of extracellular to lytic signals at 6 h in (**B**) is shown as bar graphs in (**C**). Mean ± s.d. (*n*=3 independent experiments). \*\**P*<0.01, \*\*\**P*<0.001; two-tailed Student’s *t* test. (**D** and **E**) Requirement for N-glycosylation and SP sequence for METTL9 secretion. Luciferase activities of METTL9-HiBiT WT, 2NA, and ΔSP mutants were plotted as line graphs over time (**D**). The ratio of extracellular to lytic signals at 6 h in (**D**) is shown as bar graphs in (**E**). Mean ± s.d. (*n*=3 independent experiments). \*\**P*<0.01, \*\*\**P*<0.001; two-tailed Student’s *t* test. (**F**) Expression of METTL9-HiBiT. Cell lysates expressing METTL9-FLAG WT, 2NA, and ΔSP mutants were analyzed by Western blotting with an anti-HiBiT antibody. (**G** and **H**) Immunoprecipitation of METTL9 from the medium. The culture media of cells expressing METTL9-FLAG WT and several mutants, as indicated, were immunoprecipitated with anti-FLAG antibody. These immunoprecipitates and cell lysates were analyzed by Western blotting with anti-FLAG antibody. An asterisk indicates the broad band corresponding to fully and partially N-glycosylated METTL9 (**G**). (**I** and **J**) Medium immunoprecipitation of METTL9 orthologs (I) and other histidine methyltransferases. The culture media of cells expressing FLAG-tagged METTL9 orthologs (**I**) and histidine methyltransferases (**J**) were immunoprecipitated with anti-FLAG antibody. These immunoprecipitates and cell lysates were analyzed by Western blotting with anti-FLAG antibody.

To confirm these findings, we expressed METTL9-FLAG proteins in HEK293T cells and analyzed the FLAG immunoprecipitates from the conditioned medium using Western blotting. In agreement with the HiBiT assay results, immunoprecipitation of the medium revealed the presence of wild-type METTL9, whereas the 2NA and ΔSP mutants were absent from the precipitates (Figure 3G). The secreted METTL9 was detected as a broad band at approximately 40 kDa, likely due to the maturation of N-glycosylation chains in the Golgi apparatus (* in Figure 3G). Similar experiments with the glycosylation site mutants of METTL9 showed that single site mutations at Asn-35/Thr37 or Asn-86/Ser-88 had no effect on METTL9 secretion, unlike the double mutation (Figure 3H and S3A). We also tested the possible secretion of METTL9 orthologs, as indicated in Figure 1A, and found that, in agreement with the SignalP prediction, only mammalian METTL9 (human and mouse) was secreted into the conditioned medium when expressed in HEK293T cells (Fig. 3I). On the other hand, unlike METTL9, other histidine methyltransferases known to date, including METTL18, SETD3 and CARNMT1, were not secreted in this experiment (Figure 3J). To further investigate whether METTL9 forms a disulfide-bonded dimer extracellularly, we conducted a medium immunoprecipitation assay in HEK293T cells expressing both FLAG- and HA-tagged METTL9. Under non-reducing conditions, a band that appeared to be a dimer was observed in the conditioned medium and cell lysates (Figure S3B). We also confirmed that this secretion was not mediated by extracellular vesicles (Figure S3C). Collectively, these results suggest that N-glycosylated METTL9 is secreted into the extracellular space via ER-Golgi trafficking.

### Endogenous METTL9 is secreted from neutrophil-like cells

METTL9 mRNA is widely expressed but is particularly abundant in bone marrow neutrophils [34]. This expression pattern of METTL9 led us to consider that the possible extracellular space where METTL9 functions might be in the blood flow. To test this hypothesis, we first determined the levels of METTL9 mRNA in HL-60, a human promyelocytic cell that could be differentiated into neutrophil-like cells by exposure to dimethylsulfoxide (DMSO). RT-qPCR revealed that the levels of METTL9 mRNA were approximately 6-fold higher in HL-60 cells than in HEK293T and hepatoma HepG2 cells (Figure 4A, column 1-3). Notably, a further 2-fold increase in METTL9 expression was observed upon neutrophil-like differentiation (Figure 4A, columns 3 and 4).

**Figure 4.**
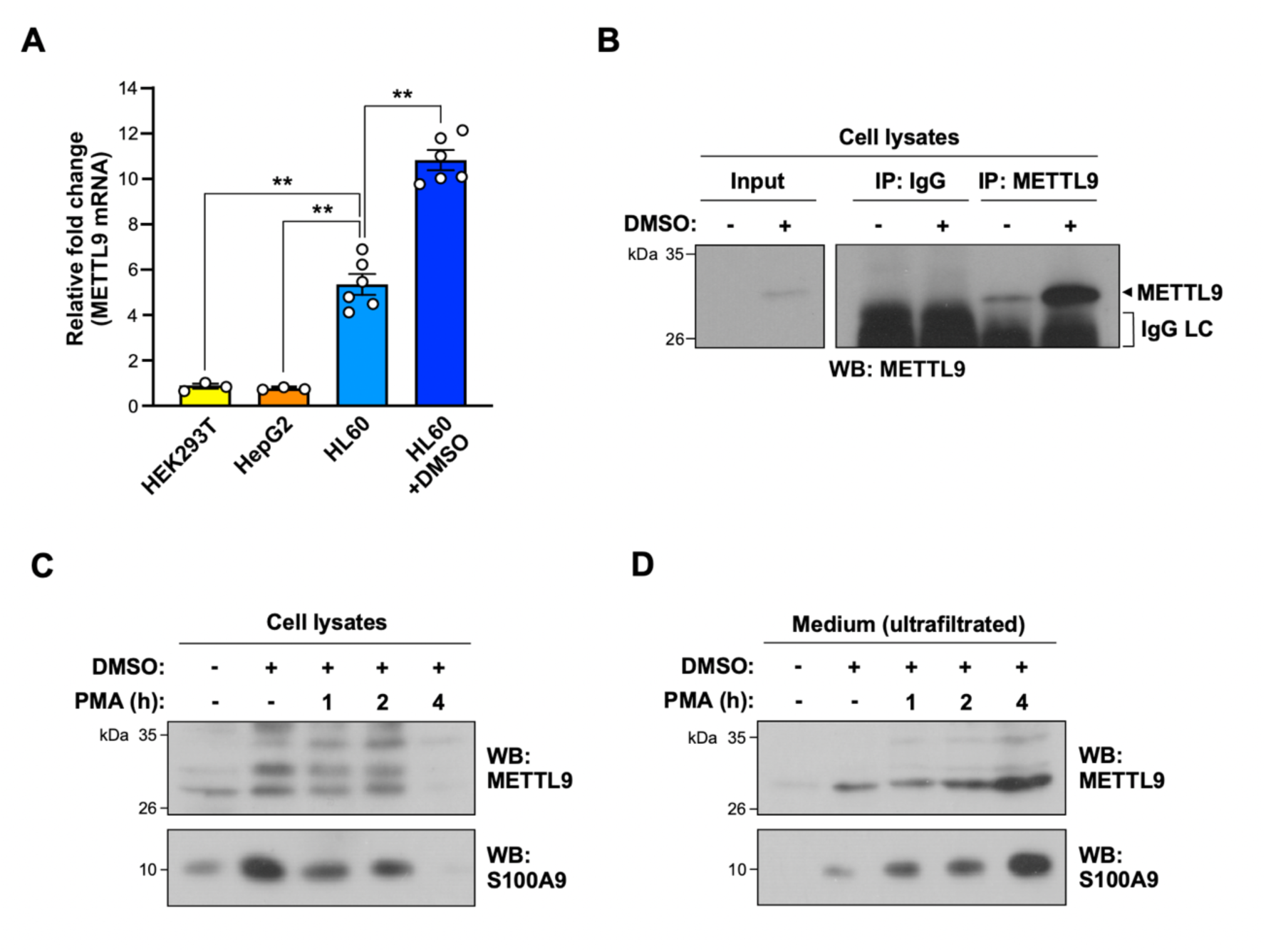
METTL9 secretion occurs in neutrophil-like cells. **(A)** mRNA expression of METTL9. Total RNAs isolated from HEK293T, HepG2, HL60, or DMSO-treated HL60 cells were analyzed using quantitative PCR and are shown as a relative ratio to the level of HEK293T cells. Mean ± s.d. (*n*=3 independent experiments). \*\**P*<0.01; two-tailed Student’s *t* test. (**B**) Endogenous METTL9 expression in HL60 cells. Lysates of HL60 cells treated with or without DMSO were immunoprecipitated with normal rabbit IgG or anti-METTL9 antibody and then analyzed by Western blotting with an anti-METTL9 antibody. An arrowhead indicates a band of endogenous METTL9. The bracket indicates the IgG light chain. (**C** and **D**) METTL9 secretion from HL60 cells. Cell lysates (**C**) and culture media (**D**) of HL60 cells treated with DMSO and/or PMA were analyzed by Western blotting with anti-METTL9 and anti-S100A9 antibodies. The culture media were concentrated by ultrafiltration before Western blotting analysis.

Next, to demonstrate the extracellular secretion of METTL9 from HL-60 cells, we first evaluated the available METTL9 antibodies for immunoprecipitation because there were no reports on the detection of endogenous METTL9. We found that the antibody purchased from Proteintech Inc. immunoprecipitated METTL9-FLAG to a similar extent as the anti-FLAG antibody (Figure S4A). Thus, we prepared lysates of undifferentiated or differentiated HL-60 cells and performed immunoprecipitation with the METTL9 antibody, followed by Western blotting. Consistent with the RT-qPCR results, METTL9 protein was detected in HL-60 cells and substantially increased after differentiation into neutrophil-like cells (Figure 4B). The specificity of immunoprecipitation was validated by reprobing with a different antibody from Atlas Antibodies (Figure S4B).

Finally, we investigated whether endogenous METTL9 is secreted extracellularly. Neutrophils release granular proteins and intracellular structures called neutrophil extracellular traps (NETs), which are composed of chromatin DNA and histones, under inflammatory conditions [35]. Therefore, we stimulated differentiated HL-60 cells with phorbol 12-myristate 13-acetate (PMA), which is known to induce neutrophil degranulation, reactive oxygen species production, and NETs formation [36]. HL-60 cells were cultured in serum-free medium with or without PMA, and cell lysates and conditioned media were collected at several time points. As shown in Figure 4C, neutrophil-like differentiation substantially induced the intracellular expression of METTL9 and S100A9, the most abundant neutrophil protein [37], whereas treatment with PMA caused a decrease in their amounts in the cells within 1 h and a complete disappearance after 4 h. In contrast, extracellular METTL9 and S100A9 were detected by neutrophil-like differentiation, and their levels increased inversely correlated with their intracellular levels after 4 h (Figure 4D). These results suggest that METTL9 is secreted from neutrophil-like differentiated HL-60 cells in response to infection and inflammatory stimuli.

### METTL9 catalyzes *N*π-methylhistidine formation of plasma proteins

Based on these results, we hypothesized that secreted METTL9 targets plasma proteins as substrates for histidine methylation. To test this hypothesis, we first assessed the enzymatic activity of the extracellular METTL9. An *in vitro* methylation assay of immunoprecipitated METTL9 from the conditioned medium demonstrated that METTL9 retains methyltransferase activity even after being secreted into the extracellular space (Figure 5A). Therefore, we investigated whether METTL9 methylates plasma proteins using an original analytical method that distinguishes between *N*π- and *N*τ-histidine methylation (Figure 5B) [18]. Before the methylation reaction, crude mouse plasma was roughly purified by removing the most abundant proteins, including albumin, IgG, and transferrin, to extend the analytical range to areas with low-abundance proteins (Figure S5A and B). Notably, the addition of wild-type METTL9 resulted in an 80-fold increase in histidine *N*π-methylation, whereas the 2GA mutant caused only a minor increase (Figure 5C, left). In contrast, no change was observed in the levels of histidine *N*τ-methylation in the presence or absence of METTL9 (Figure 5C, right). Moreover, column-purified high-abundance serum proteins were not methylated by METTL9 (data not shown). These data suggest that plasma proteins other than albumin, IgG, and transferrin could be targets of METTL9-induced methylation *in vitro*.

**Figure 5.**
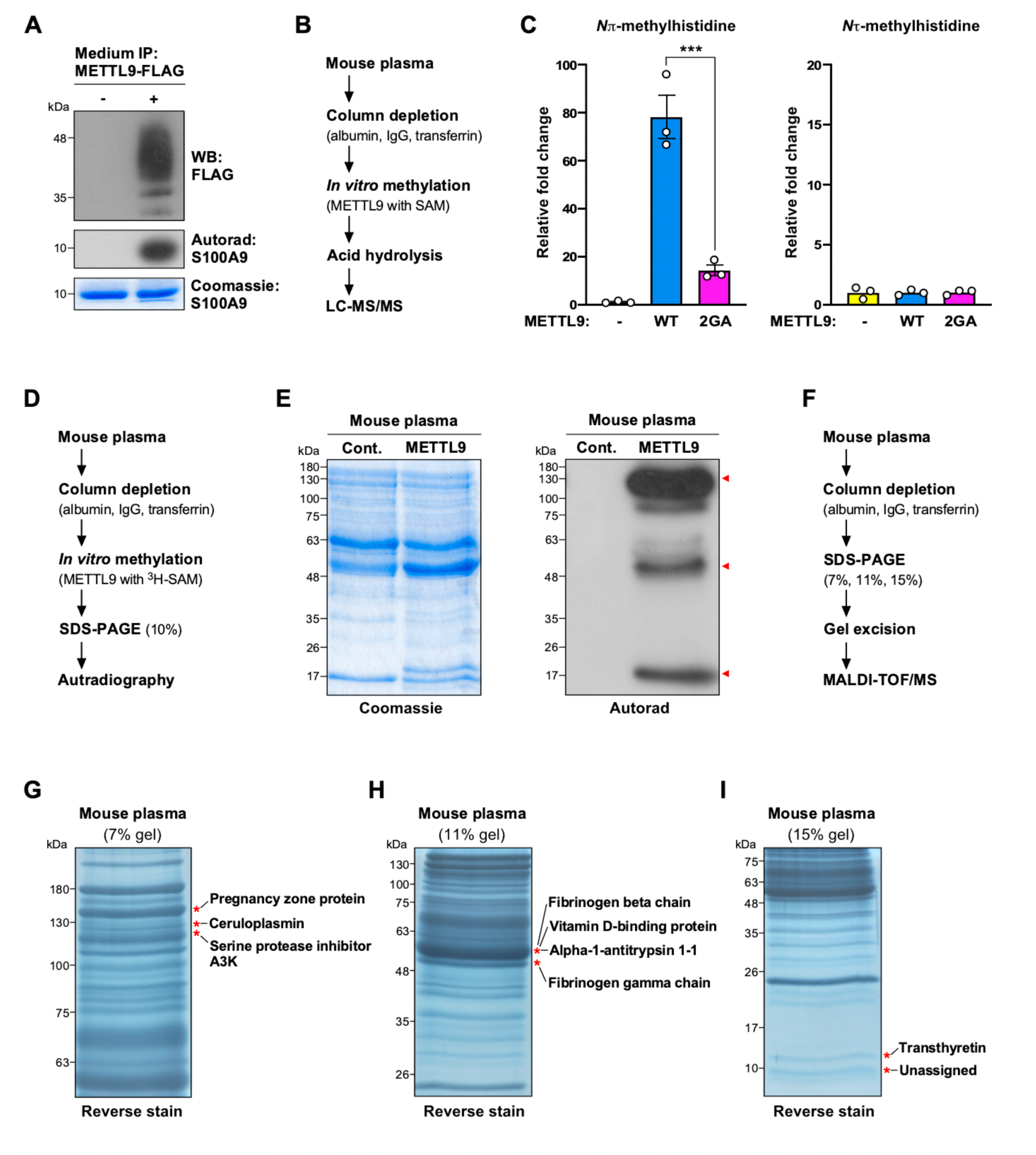
METTL9 methylates plasma proteins *in vitro*. **(A)** Secreted METTL9 methylates S100A9. Immunoprecipitated METTL9-FLAG from the culture medium was incubated with recombinant S100A9 and ^3^H-labeled SAM. METTL9-FLAG and S100A9 in the reaction were detected by Western blotting (*Top*) and Coomassie blue staining (*Bottom*), respectively. The incorporation of [^3^H]-methyl into S100A9 was visualized by autoradiography (*Middle*). (**B**) Schematic representation of the strategy used to measure the levels of histidine methylation in plasma proteins after *the in vitro* methylation assay. (**C**) *N*π-(left) and *N*τ- (right) methylhistidine levels in plasma proteins after *in vitro* methylation assay. Column-purified plasma was incubated with recombinant METTL9 WT or 2GA mutant in the presence of SAM, followed by acid hydrolysis. The methylhistidine amounts were determined by LC-MS/MS and are shown as a relative ratio to the no-enzyme control. Mean ± s.d. (*n*=3 independent experiments). \*\**P*<0.01; two-tailed Student’s *t* test. (**D**) Schematic representation of the strategy for detecting histidine methylation in plasma proteins after *an in vitro* methylation assay. (**E**) *In vitro* methylation of plasma proteins by METTL9. Column-purified plasma was incubated with or without recombinant METTL9 WT in the presence of ^3^H-labeled SAM, followed by SDS-PAGE. Total proteins in the reaction were detected using Coomassie blue (*Left*). The incorporation of [^3^H]-methyl into plasma proteins was visualized using autoradiography (*Right*). Red arrowheads indicate major methylation bands. (**F**) Schematic representation of the strategy to identify candidate proteins for histidine methylation in plasma based on the results in (**E**). (**G**-**I**) Identification of candidate(s) for METTL9-induced methylation in plasma proteins. Column-purified plasma was separated by 7% (**G**), 11% (**H**), and 15% (**I**) SDS-PAGE, followed by reverse staining. Asterisks indicate the bands analyzed by MALDI-TOF/MS. The names of the identified proteins are listed.

Thus, we next sought to identify METTL9 substrates for methylation by detecting the incorporation of the [^3^H]-methyl group of [^3^H]-*S*-adenosylmethionine into the mouse plasma proteins (Figure 5D). After 10% SDS-PAGE separation, several methylation signals were obtained by fluorography around the molecular weight markers at 130, 48, and 17 kDa (Figure 5E, right). To identify the proteins corresponding to the bands detected at these heights, column-purified mouse plasma was finely separated by 7%, 11%, and 15% SDS-PAGE, followed by reverse staining. The bands corresponding to the methylation signal heights on each gel were excised and subjected to MALDI-TOF/MS analysis (Figure 5F). As shown in Figure 5G-I and Figure S5C-J, eight plasma proteins were identified as possible targets for METTL9-induced methylation: pregnancy zone protein, ceruloplasmin, serine protease inhibitor A3K, fibrinogen beta chain, vitamin D-binding protein, alpha-1-antitrypsin 1-1, fibrinogen gamma chain, and transthyretin. Among them, we focused on ceruloplasmin (hereafter referred to as CP) and transthyretin (hereafter referred to as TTR) because both possess the “His-x-His motif” (H-x-H, where x is preferably a small amino acid), which is the consensus sequence for METTL9-dependent methylation (Figure S5K and L). CP is a copper transporter in the bloodstream and has ferroxidase activity, oxidizing Fe^2+^ to Fe^3+^ [38], while TTR acts as a carrier protein for thyroid hormones, such as tetraiodothyroxine (T_4_), and retinol (vitamin A) in blood plasma and cerebrospinal fluid [39].

### METTL9 directly methylates transthyretin and ceruloplasmin

Since TTR is a much smaller protein than CP (147 aa vs. 1061 aa) and has only one H-x-H motif, we first addressed the possibility of methylation in TTR. An *in vitro* METTL9 methylation assay was performed using recombinant mouse TTRΔSP lacking the N-terminal signal peptide (1-20 aa) in the presence of SAM, and the methyl amino acid content of TTR was analyzed using LC-MS/MS (Figure 6A). As shown in Figure 6B, METTL9 selectively catalyzed the formation of *N*π-methylhistidine in TTR *in vitro*. Importantly, similar results were obtained when the human orthologs of TTR and METTL9 proteins were used in *the in vitro* methylation assay (Figure S6A and B), implying that TTR is a *bona fide* substrate for METTL9, at least in mammals.

**Figure 6.**
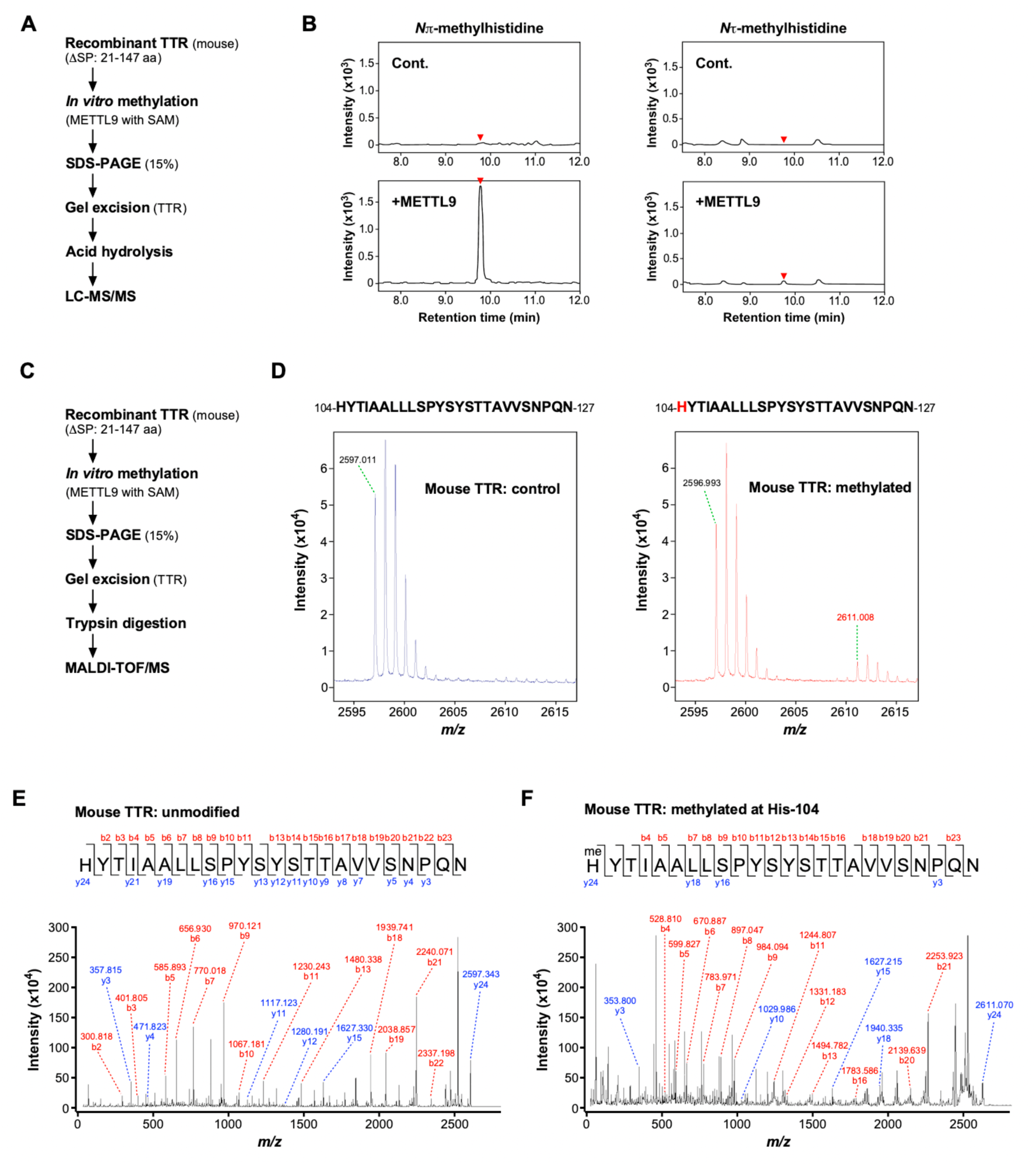
METTL9 methylates transthyretin at His-124. **(A)** Schematic representation of the strategy used to measure the levels of histidine methylation in mouse TTR after *the in vitro* methylation assay. (**B**) METTL9 catalyzes *N*π-methylhistidine formation in mouse TTR *in vitro*. Recombinant TTR was incubated with recombinant METTL9 WT in the presence of *S*-adenosylmethionine (SAM). After SDS-PAGE, the methylhistidine content of TTR was determined by LC-MS/MS and is shown as chromatograms. The red arrowheads indicate the retention times of *N*π- and *N*τ-methylhistidine as defined by the standard. (**C**) Schematic representation of the strategy to identify methylation sites of mouse TTR after *in vitro* methylation assay. (**D**) Tryptic peptides of mouse TTR contain monomethylated amino acids. *In vitro* methylation reactions were resolved using SDS-PAGE, followed by reverse staining. The portion of the gel corresponding to TTR was digested with trypsin and analyzed using MALDI-TOF/MS. Unmodified (*Left*) and monomethylated (*Right*) peptide sequences from mouse TTR are shown. The METTL9-mediated methylation site at His-104 is denoted in red. (**E** and **F**) MS/MS fragmentation spectra showing monomethylation of mouse TTR at His-104. The 24-amino acid peptide sequence and b-series (red) and y-series (blue) peptide fragment ions for unmodified (**E**) and methylated (**F**) peptide products are shown.

Next, to determine which histidine residues are methylated in TTR, we explored methylated TTR-derived peptides using MALDI-TOF MS analysis after an *in vitro* methylation assay (Figure 6C). A peptide containing a monomethylated amino acid from residues 104–127 of TTRΔSP was identified by trypsin digestion (Figure 6D). Furthermore, we investigated the specific methyl group localization using MS/MS analysis. As expected, it was confirmed that His-104 overlapping with the second histidine residue in the HxH motif is a METTL9-dependent monomethylation site in mouse TTR (Figure 6E and F). Likewise, we identified the methylation site of human TTR at His-90 (Figure S6C-F).

We then examined whether METTL9 methylates CP *in vitro*. Because full-length CP protein could not be synthesized in *E. coli*, FLAG-tagged CP was overexpressed in HEK293T cells, and the immunopurified product was used as the substrate (Figure S7A). LC-MS/MS analysis demonstrated that METTL9 introduces *N*π-methylhistidine in CP as well as TTR (Figure S7B). Next, we prepared bacterially expressed N-terminal (NT: 20-205 aa) and C-terminal (CT: 725-1061) regions of CP proteins, which contain two and five H-x-H motifs, respectively, and determined which histidine residues were methylated by METTL9 (Figure S7C and D). In the CP-NT region, both His-122 and His-181 were identified as target sites for METTL9-induced methylation (Figure S7E-H); however, only His-832 was found to be methylated in the CP-CT region (Figure S7I and J). Collectively, these results indicate that at least two plasma proteins, TTR and CP, are substrates for METTL9-induced methylation.

### Histidine methylation of transthyretin decreases the binding affinity to zinc

Finally, to clarify the significance of plasma protein methylation by METTL9, we investigated the impact of TTR histidine methylation on its molecular functions. TTR is a β-sheet-rich protein that exists primarily as a homotetramer in blood plasma and cerebrospinal fluid [39]. As shown in Figure 7A, the positions of methylated histidine residues in humans and mice are not completely identical, but they are in close proximity. In particular, since the methylation site of human TTR at His-90 is located immediately adjacent to the β-strand structure (referred to as β-strand F), which acts as a dimerization surface interacting with the β-strand H of another subunit by hydrogen bonds (Figure 7A and B) [40], we investigated whether His-90 contributes to the dimer formation of TTR. To this end, we performed chemical crosslinking assays using culture media and whole cell lysates of HEK293T cells expressing either TTR wild type, the HxH motif mutants (H88A and H90A), or the known monomeric mutant (F87E) [41]. However, unlike the F87E mutant, the H88A and H90A mutants formed dimers to the same extent as the wild-type (Figure 7C and D), suggesting that the chemical properties of His-88 and His-90 residues are not involved in TTR dimer formation, at least in this experiment.

**Figure 7.**
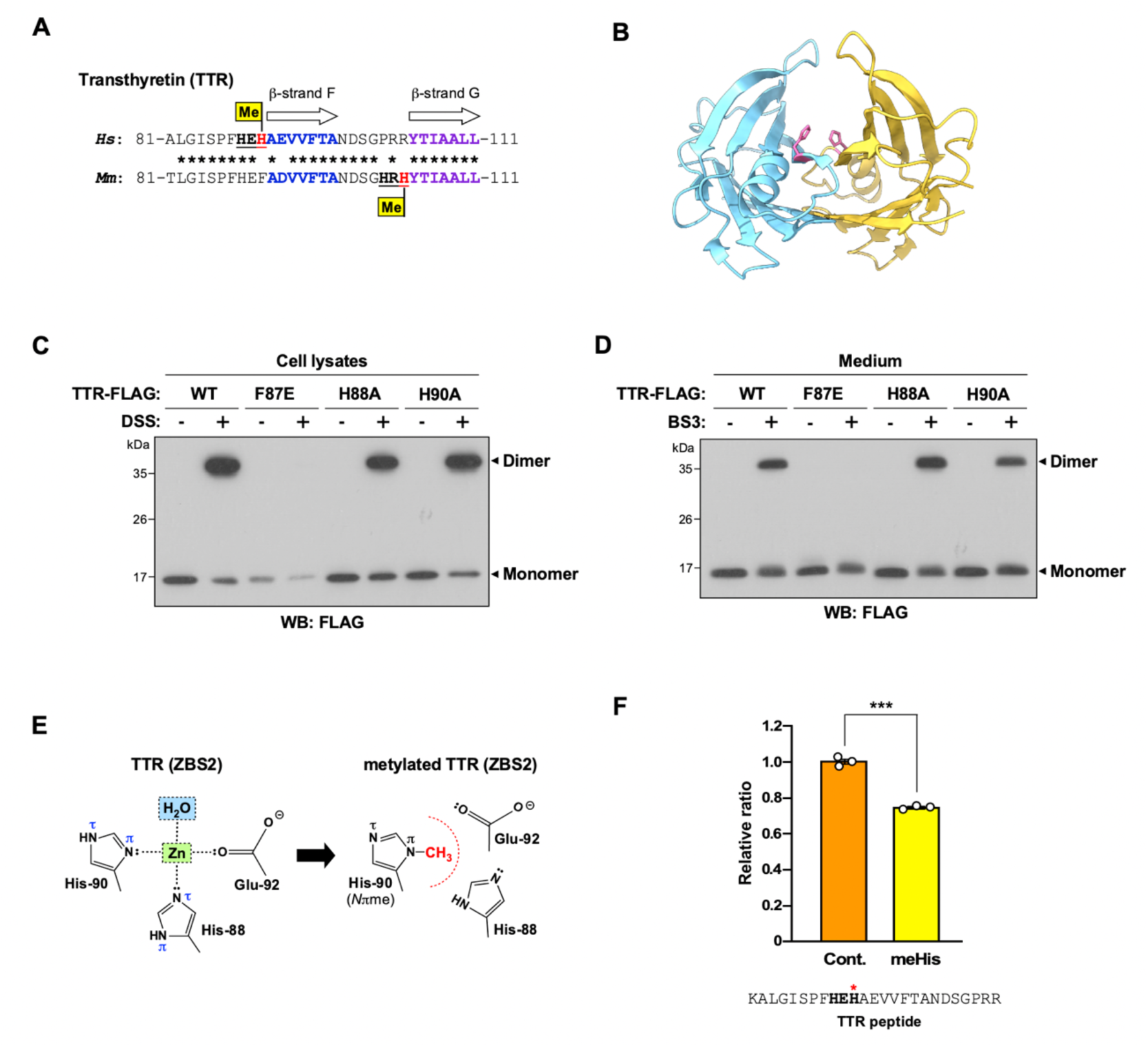
METTL9-mediated methylation of TTR decreases the binding affinity to zinc. **(A)** Comparison of amino acid sequences around methylation sites in human and mouse TTR. *Hs*, *Homo sapiens*; *Mm*, *Mus musculus*. The HxH motifs are shown in bold and underlined. The methylation sites at His-90 (*Hs*) and His-104 (*Mm*) are denoted in red and indicated by a flag. β-strands F and G are shown in bold blue and bold purple, respectively. Conserved sequences are indicated by asterisks (*). (**B**) Cartoon representation of the human TTR dimer (PDB code: 319P). The α-helices are shown as ribbons, β-strands as arrows, and loops as tubes. The His-90 side chain is shown in pink. (**C**) Chemical crosslinking of intracellular TTR. HEK293T cells expressing TTR-FLAG WT or F87E, H88A, or H90A mutants were treated with DSS crosslinker and cell lysates were analyzed by Western blotting with anti-FLAG antibody. (**D**) Chemical crosslinking of extracellular TTR. The culture media of cells expressing TTR-FLAG WT or F87E, H88A, or H90A mutants were treated with BS3 crosslinker and then analyzed by Western blotting with anti-FLAG antibody. (**E**) Close-up views of Zn^2+^ binding site (ZBS2) and schematic diagrams of Zn^2+^ coordination spheres. ZBS2 is composed of His-88, His-90, Glu-92, and one water molecule (*Left*). *N*π-methylation of His-90 blocks the coordination of Zn^2+^ (*Right*). (**F**) *N*π-methylation of His-90 in the TTR peptide attenuates its binding to zinc. A synthetic human TTR peptide containing either no (control) or *N*π-methylhistidine (meHis) residue was passed through a zinc-chelating column, and the bound peptide was measured using LC–MS/MS. The ratio of the bound peptide concentrations of control to meHis is shown as bar graphs. The peptide sequence used in this assay is shown, and the methylated histidine corresponding to His-90 of TTR is indicated by a red asterisk (*). Mean ± SD (n = 3 independent experiments). \*\*\**P*<0.001; two-tailed Student’s *t* test.

Meanwhile, TTR is also known as a zinc-binding protein, possessing three zinc-binding sites (ZBSs) within each monomer, and one of these, the HxHxE motif composed of His-88, His-90, and Glu-92, is referred to as ZBS2 [42]. Importantly, in ZBS2, Zn^2+^ is bound to the imidazole nitrogens of His-88 (*N*τ atom) and His-90 (*N*π atom), the carboxylic acid group of the Glu-92 side chain (Oδ1 atom), and a water molecule (Figure 7E, left). Considering that His-90 undergoes *N*π-methylation by METTL9, this modification is likely to inhibit the coordination bonding with Zn^2+^ (Figure 7E, right). To test this possibility, we synthesized a TTR-derived peptide containing the ZBS2 motif (residues 80–114) and a methylated version of this peptide at His-90 and compared their zinc-binding activities using metal-chelate chromatography on iminodiacetic acid equilibrated with zinc. Interestingly, only one histidine *N*π-methylation of the TTR peptide resulted in a 25% reduction in zinc binding, suggesting that METTL9-mediated methylation decreases the zinc-binding ability of TTR as with the case of S100A9 (Figure 7F) [18].

## Discussion

Here, we discovered that histidine *N*π-methyltransferase METTL9 is secreted into the extracellular space. This is the first report of a secreted methyltransferase and the second example of a secreted post-translational modification enzyme, following secreted kinases such as Fam20C. Consistent with Fam20C, METTL9 has an SP sequence, localizes to the ER-Golgi compartments, and undergoes N-linked glycosylation. Furthermore, we demonstrate that METTL9 forms disulfide bonds during dimer formation. Given that disulfide bonds are a common feature of secreted plasma proteins that stabilize the structure and protect the thiol group from over-oxidation, the disulfide dimer of METTL9 appears to allow robust dimer formation even in harsh extracellular environments, where high stability is usually required for functional activity. This supports our hypothesis that METTL9 acts as an endocrine methyltransferase in the bloodstream.

Why is METTL9 secreted by neutrophils? In our previous report, we demonstrated that METTL9 directly methylates the proinflammatory protein S100A9, which is highly expressed accounting for 40% of the cytoplasmic proteins in neutrophils, and this methylation attenuates the zinc-binding affinity of S100A9 *in vitro* [18]. Supporting our findings, Cao *et al*. have provided *in vivo* evidence for METTL9-catalyzed methylation of S100A9 in mice and revealed its physiological role in response to *Staphylococcus aureus* infection [43]. They proposed a four-step model for the METTL9-S100A9 methylation axis in the regulation of the host’s antibacterial responses: (i) METTL9-induced S100A9 methylation occurs in neutrophils at steady state; (ii) Upon *S. aureus* infection, S100A9 methylation rapidly decreases (the acute stage); (iii) Hypomethylated S100A8/A9 sequesters zinc ions from *S. aureus* to restrict its growth; and (iv) After the clearance of bacteria, S100A9 methylation gradually increases to prevent excessive zinc chelation and promote tissue repair (the resolving stage) [43]. In light of our findings, it is likely that in the resolving stage (iv), secreted METTL9 actively blocks the chelating ability of zinc by methylating extracellular S100A9, which is already secreted and present at the inflammatory site. In addition, it remains unclear why the methylation levels of S100A9 decrease in neutrophils upon *S. aureus* infection (ii). However, considering that no histidine demethylase has yet been identified, the extracellular release of METTL9 might consequently be a factor in lowering the methylation of intracellular S100A9.

In addition to its role in transporting thyroid hormones and retinol in blood plasma and cerebrospinal fluid, TTR is recognized as a primary amyloidogenic protein [44]. It forms extracellular amyloid deposits in nerves, ligaments, and the heart and is associated with two forms of systemic amyloidosis: hereditary ATTR (ATTRv) amyloidosis induced by variant TTR and aging-related wild-type ATTR (ATTRwt) amyloidosis. Although the destabilization of TTR tetramers is widely believed to be a critical step in amyloid fibril formation, the precise molecular mechanism remains unknown. However, given that a reduction in pH induces the dissociation of the tetrameric form of TTR *in vitro* [45], histidine methylation has emerged as a potential mechanism of amyloid formation, as described below. Specifically, the imidazole ring of histidine residues can readily transition between its protonated (positively charged) and unprotonated (neutral) forms within the physiological pH range; however, this chemical property as a local pH sensor can be prevented by histidine methylation, thereby affecting the stability of the TTR tetramer. Supporting this notion, continuum electrostatic analysis of human TTR has shown that in the pH range of 3.5-7.0, His-88 and His-90 as well as Tyr-116 are the most destabilizing residues upon dimer formation [46]. Furthermore, neutron diffraction studies of human TTR have revealed that the protonation state of the histidine residue is closely related to tetramer stability, and the double protonation of His-88 by acidification is predicted to break the hydrogen-bond network and cause the destabilization of the TTR tetramer [47, 48]. Notably, these studies also found a weak hydrogen bond between His-90 and Glu-92, implying that TTR methylation at His-90 may affect the hydrogen bond in the TTR tetramer. Based on this hypothesis, a low-grade, chronic, and systemic inflammatory state coupled with aging (inflammaging) might be a cause of the pathogenic mechanism of aging-associated ATTRwt amyloidosis, where secreted METTL9 from neutrophils induces TTR methylation, thus resulting in the dissociation of the tetrameric structure into an altered monomeric structure, which eventually leads to amyloid fibrils.

Meanwhile, TTR has been shown to possess proteolytic activity in addition to its well-known transport capabilities [39]. It was described as a cryptic peptidase that cleaves several natural substrates, such as apolipoprotein A-I and β-amyloid, each of which is involved in atherosclerosis and Alzheimer’s disease, respectively [49, 50]. Notably, Liz MA *et al*. have reported that the peptidase activity of human TTR requires Zn^2+^-binding to the catalytically important residues His-88, His-90, and Glu-92, establishing a Zn^2+^-chelating pattern known as HxHxE [51]. In fact, they showed that replacing even one of these side chains with alanine almost abolishes peptidase activity using fluorogenic peptides for TTR cleavage. Given our results that the reduced Zn^2+^-binding capacity of the methylated TTR peptide at His-90 compared to the unmodified peptide, it is possible that METT9-induced TTR methylation may be involved in the inhibitory regulation of the metallopeptidase activity of TTR in plasma.

In this study, we focused on blood as an extracellular fluid where METTL9 functions; however, the extracellular domain of transmembrane proteins and fibrous proteins in the extracellular matrix (ECM) could be possible targets for methylation. For example, integrins, which are cell adhesion molecules that mediate the adhesion of cells to the ECM and other cells, form heterodimers composed of two distinct subunits, α and β [52]. Intriguingly, of the total 18 α subunits and 8 β subunits in humans, 10 (1, 2, 3, 4, 7, 9, 10, E, L, M) for α and 3 (4, 7, 8) for β have the HxH motif, which is the consensus sequence for METTL9-induced methylation. Among them, αLβ2 integrin (also called LFA-1) is widely expressed on leukocytes, including neutrophils, and is involved in leukocyte migration to inflammatory sites, suggesting that secreted METTL9 from neutrophils may modulate their adhesion to the vascular endothelium or extravasation. Moreover, it has been reported that elevated METTL9 expression contributes to the migration and invasive potential of scirrhous gastric carcinoma [53]. In view of the crucial roles of cell adhesion molecules, such as integrins, in cancer metastasis, secretory METTL9 may be implicated in these processes by methylating the extracellular domain of transmembrane proteins and fibrous proteins in the ECM.

While we demonstrated that METTL9 is highly expressed in neutrophils, a recent study by Codino *et al*. found abundant expression of METTL9 in the mouse and human brains during development [54]. Using three different mouse embryonic stem cell lines (a Mettl9 knockout, an inducible METTL9 Degron, and a catalytically inactive Mettl9 knock-in) and *Xenopus laevis* embryos, the authors revealed that METTL9 plays a conserved role in sustaining vertebrate neurogenesis. Interestingly, they showed that METTL9 modulates the secretory pathway in the Golgi apparatus independent of its catalytic activity by binding to regulatory factors involved in cellular transport, endocytosis, and Golgi integrity. Combining their findings with our data, METTL9 may possess a dual function both inside and outside the cell. Specifically, it maintains the secretory system in the Golgi apparatus without methylating key regulators, while also being secreted into the extracellular space to function as a methylating enzyme. Also noteworthy is that, in our experiments, the METTL9 (2GA) mutant, which lacks methylation activity, was not secreted extracellularly despite undergoing N-glycosylation to the same extent as the wild type (data not shown). Considering that METTL9 (2GA) is unable to bind to SAM due to mutation, alterations in SAM levels within the Golgi apparatus may determine whether METTL9 is secreted extracellularly or retained intracellularly.

In conclusion, the discovery of methyltransferase secretion suggests the potential for the existence of diverse extracellular PTMs, extending beyond phosphorylation and histidine methylation, which in turn might serve as post-secretory activity regulation of endocrine proteins, such as insulin and growth hormones, in the cardiovascular system.

## Experimental procedures

### Mammalian cell culture

The human embryonic kidney cell line expressing SV40 large T antigen, HEK293T (RCB2202), was provided by the RIKEN BRC through the National BioResource Project of the MEXT, Japan. HEK293T cells were cultured in DMEM (Nacalai tesque #08456-36) supplemented with 10% fetal bovine serum (Sigma-Aldrich #F7524) and 1% penicillin-streptomycin solution (Nacalai tesque #26253-84) at 37°C in a humidified atmosphere with 5% CO_2_. The human promyelocytic cell line HL60 (JCRB0085) was provided by the JCRB Cell Bank, Japan. HL60 cells were cultured in suspension in RPMI1640 (Nacalai tesque #05176-25) supplemented with 20% fetal bovine serum (Sigma-Aldrich #F7524) and 1% Penicillin-Streptomycin solution (Nacalai tesque #26253-84) at 37°C in a humidified atmosphere with 5% CO_2_. Neutrophil-like differentiation was induced by seeding the cells at a concentration of 0.5 × 10^6^ cells/mL in the presence of DMSO at a final concentration of 1.25% (v/v) for 5 days. For treatment with phorbol 12-myristate 13-acetate (PMA; Selleck #S7791), the cells were centrifuged at 2,000 rpm for 2 min and resuspended in serum-free RPMI1640 medium in the presence or absence of PMA (100 nM), followed by incubation for up to 4 h. Cell concentration and viability were determined by trypan blue dye exclusion using a TC10 automated cell counter (Bio-Rad #145-0001).

### Mice

C57BL/6J mice were obtained from Clea Japan (Tokyo, Japan) and housed in a specific pathogen-free barrier facility with a 12-h light/dark cycle and provided *ad libitum* access to standard chow (MF diet, ORIENTAL YEAST) and water. Animal experiments were performed in a humane manner and were approved by the local ethics committee of the University of Tsukuba. Experiments were performed in accordance with the Regulation of Animal Experiments of the University of Tsukuba and the Fundamental Guidelines for Proper Conduct of Animal Experiments and Related Activities in Academic Research Institutions under the jurisdiction of the Ministry of Education, Culture, Sports, Science, and Technology, Japan.

### Gene cloning and plasmid construction

The coding sequences of *mettl9* (human), *mettl18* (human), *setd3* (human), and *carnmt1* (mouse), and *transthyretin* (mouse and human) were amplified using PrimeSTAR MAX DNA polymerase (Takara Bio #R045A) from cDNAs derived from total RNA of HEK293T cells, HepG2 cells, or mouse liver. The coding sequences of *the mettl9* orthologues of *Danio rerio* (F1R335) and *Drosophila melanogaster* (Q9W1H1) were artificially synthesized by Eurofins Genomics. Point mutations were generated by Q5 Site-Directed Mutagenesis Kit (New England Biolabs #E0552S) and specific primers. The coding sequences were subcloned into the pcDNA3 vector with a C-terminal FLAG-tag (*mettl9*, *mett18*, *setd3*, *carnmt1*, *transthyretin*) or HiBiT-tag (*mettl9*) or pMAL-c6T vector (*transthyretin*) using the In-Fusion Snap Assembly Cloning Kit (Takara Bio #638948). All constructs were verified by sequencing using the SupreDye Cycle Sequencing Kit (EdgeBio #060002) on a 3130*xl* Genetic Analyzer (Applied Biosystems). The other constructs including pcDNA3/METTL9-HA (mouse), pGEX-6P/METTL9 (mouse and human), and pGEX-6P/S100A9 (mouse and human) were described previously [18].

### Transfection

Transfection of plasmid DNA was performed using GeneJuice Transfection Reagent (Sigma-Aldrich #70967) at a ratio of 2 μL GeneJuice / 1 μg DNA on 60% confluent cells, according to the manufacturer’s instructions. Cells were grown for 24-36 h post transfection before treatment or collection.

### Preparation of whole-cell lysates

Cells were washed twice with PBS and lysed for 15 min on ice in a cell lysis buffer containing 20 mM HEPES-KOH (pH 7.9), 150 mM NaCl, 0.1% Triton X-100, 0.1 mM EDTA, and 1 × protease inhibitor cocktail (Nacalai Tesque #25955). The homogenate was centrifuged at 19,000 × *g* for 10 min at 4°C, and the supernatant comprising the whole cell lysate was mixed with an equal volume of 2 × SDS sample buffer containing 12% β-mercaptoethanol and boiled for 5 min. Under non-reducing conditions, whole cell lysates were mixed with an equal volume of 2 × SDS sample buffer containing 10 mM iodoacetamide and incubated at 37°C for 1 h. The protein concentrations in the whole-cell lysates were determined using the Quick Start Bradford Protein assay kit (Promega #5000204) with bovine γ-globulin as the external standard.

### Western blotting

Whole-cell lysates, immunoprecipitates, and culture media were separated by SDS-polyacrylamide gel electrophoresis and transferred to a 0.45 μM PVDF membrane (Millipore #IPVH00010) using a Trans-Blot Turbo Transfer System (Bio-Rad #1704150). After blocking with 0.3% skim milk in TBS supplemented with 0.1% Tween-20 (TBST) for 1 h, the membrane was incubated for 1 h at room temperature or overnight at 4°C with primary antibodies in 0.3% skim milk in TBS-T, followed by washing three times with TBS-T. The membrane was then probed with horseradish peroxidase-conjugated secondary antibodies for 30 min at room temperature, followed by washing three times with TBS-T. To visualize the protein bands, the membrane was developed with Clarity Western ECL Substrate (Bio-Rad #1705061) and exposed to Super RX X-ray film (Fujifilm #47410 26617).

### Immunoprecipitation

Whole-cell lysates prepared as described above or culture media cleared by centrifugation (800 × g for 1 min at 4°C) were incubated with either ANTI-FLAG M2 Affinity Gel (Millipore #A2220) or antibody-conjugated Protein G Sepharose (Cytiva #17061801) at 4°C, rotating for 3 h to overnight. After washing three times with ice-cold cell lysis buffer (whole-cell lysates) or PBS (culture media), the beads were used for Western blot analysis by adding SDS sample buffer. For *the in vitro* methylation assay, immunoprecipitated METTL9-FLAG beads from whole-cell lysates and culture media were used as methyltransferases.

### Identification of METTL9-interacting protein by MALDI-TOF/MS

To identify METTL9-interacting proteins using co-immunoprecipitation assay, FLAG-immunoprecipitates were separated by SDS-PAGE and stained with Coomassie Brilliant Blue R-250. A specific band observed in METTL9-FLAG WT but not in the 2NA mutant at approximately 100 kDa was excised and digested in gel with trypsin after reductive alkylation treatment with DTT and iodoacetamide. After 24 h of incubation at 37 °C in incubation buffer (50 mM NH_4_HCO_3_ and 1 mM CaCl_2_), the tryptic peptides were eluted from the polyacrylamide gel by sonication in acetonitrile containing 0.1% formic acid. The peptides were desalted with ZipTip μC_18_ (Millipore, #ZTC18M096). A saturated solution (10 mg/ml) of *R*-cyano-4-hydroxycinnamic acid (Shimadzu GLC) in 0.1% trifluoroacetic acid and 50% acetonitrile was used as the MALDI matrix. MS analysis was conducted using ultrafleXtreme NTA (Bruker), and the acquired data were processed by FlexAnalysis (Bruker) and BioTools (Bruker) software.

### Expression and purification of recombinant proteins

GST-fused METTL9 (mouse and human), S100A9 (mouse), and TTR (mouse and human) were expressed in *Escherichia coli* and batch-purified using an affinity chromatography resin. In brief, the pGEX-6P constructs encoding the proteins described above were transformed into chemically competent BL21-CodonPlus (DE3)-RIL strain (Agilent #230240). The bacteria were cultured in LB medium containing ampicillin (100 μg/mL) until the OD600 reached 0.8, and protein expression was induced by adding 0.5 mM IPTG (Nacalai tesque #06289-67) at 25°C for 3 h. Cells were collected by centrifugation (3,000 rpm for 10 min) and resuspended in PBS supplemented with lysozyme (1 mg/mL, Nacalai tesque #19499-04), followed by incubation on ice for 10 min. After adding 0. 5% NP-40, 1 mM DTT, and 1 × protease inhibitor cocktail, the suspended cells were sonicated at 20 kHz (amplitude 40; Qsonica Q55) for 3 × 20 s, interspaced by more than 30 s on ice, and then rotated for 15 min at 4°C. Cell debris and insoluble fractions were removed by centrifugation at 19,000 × *g* for 10 min at 4°C and the supernatants were mixed with 50 μL bed volume of equilibrated Glutathione Sepharose 4B (Cytiva, #17513201) at 4°C for at least 2 h. The fusion protein-bound beads were washed three times with ice-cold PBS and once with cleavage buffer [50 mM Tris-HCl (pH 7.0), 150 mM NaCl, 1 mM EDTA, 1 mM DTT]. The beads were then incubated with 16 units of PreScission Protease (Cytiva #27084301) in 100 μL of cleavage buffer at 4°C overnight. The eluates were collected by centrifugation at 500 × *g* for 5 min, aliquoted, stored at -80°C, and used for *in vitro* methylation assays.

Maltose-binding protein (MBP)-fused mouse CP fragments were expressed in *Escherichia coli* and batch-purified using affinity chromatography resin. Briefly, the pMAL-c6T constructs encoding CP-NT (20-205 aa) and CP-CT (725-1061 aa) were transformed into chemically competent Rosetta 2 (DE3) strain (Sigma-Aldrich #71397). Thereafter, the cell lysates were prepared using the same procedure as described above and mixed with 50 μL bed volume of equilibrated Amylose resin (NEB, #8021S) at 4°C for at least 2 h. The fusion protein-bound beads were washed three times with ice-cold PBS and used for *in vitro* methylation assays.

### Treatment with deglycosylating enzyme

Peptide:*N*-glycanase F from *Escherichia coli* (Glycopeptidase F; Takara Bio #4450) was used for deglycosylation of METTL9-FLAG, according to the manufacturer’s instructions. Briefly, immunoprecipitated METTL9-FLAG from HEK293T cells was eluted with 0.1 M glycine-HCl (pH 2.5), followed by dialysis with a Slide-A-Lyzer dialysis cassette (3.5 kDa MWCO). Immunopurified METTL9-FLAG (2.5 μL) was mixed with equal volumes of denaturing buffer [1% SDS, 1 M Tris-HCl (pH 8.6), 0.2 M 2-mercaptoethanol] and then heat denatured at 100°C for 3 min. After adding 5% NP-40 (5 μL) and purified water (13 μL), the mixture was incubated in the presence or absence of Glycopeptidase F (1 mU) at 37°C for 20 h. The reaction mixture was separated using SDS-PAGE and analyzed by Western blotting.

### *In vitro* methylation assay with radioisotope

*In vitro* methylation reactions against S100A9 were performed using immunopurified METTL9-FLAG WT, 2NA, or 2GA from HEK293T cells, together with affinity-purified recombinant S100A9 (3 μg) in the presence of *S*-[methyl-^3^H] adenosyl-L-methionine (1 μCi; Revvity #NET155V) in PBS. After incubation at 30°C for 1 h, the reactions were resolved by SDS-PAGE, and the gels were stained with Coomassie blue, followed by soaking in Amplify fluorographic reagent (GE Healthcare) for 20 min. The gels were dried under vacuum at 80°C for 1 h and then exposed to Amersham Hyperfilm ECL (GE Healthcare) at -80°C for 2 days.

### HiBiT extracellular and lytic detection assays

Nano-Glo HiBiT extracellular detection assay was performed according to the manufacturer’s protocols with some modifications. Briefly, HEK293T cells seeded in a collagen-coated 12-well plate (IWAKI) were transfected with METTL9-HiBiT expression plasmids (0.5 μg/well) and incubated for 24 h after the transfection. After washing twice with PBS, the cells were incubated with 1mL/well DMEM supplemented with 10% fetal bovine serum in the presence or absence of 1 μM brefeldin A for 0, 2, 4, or 6 h. Fifty μL of culture media at each time point were transferred into white flat bottom 96-well plate and mixed with equal volume of the HiBiT nonlytic reagent containing the Nano-Glo HiBiT extracellular buffer, the cell-impermeable LgBiT protein, and the furimazine substrate in a ratio of 100: 1: 2. After incubation in a light-shielded environment at room temperature for 10 min, luminescence was measured using a Centro LB960 plate luminometer (Berthold). Additionally, to assess intracellular expression of METTL9-HiBiT, HEK293T cells post collecting the culture media were washed twice with PBS and lysed for 15 min on ice in the HiBiT lytic reagent (400 μL) containing the Nano-Glo HiBiT lytic buffer, the LgBiT protein, and the furimazine substrate in a ratio of 100: 1: 2. The homogenate was centrifuged at 19,000 × *g* for 10 min, and the luminescence of the supernatant was measured using a Centro LB960 plate luminometer (Berthold).

### Extracellular vesicle isolation from culture medium

An Exosome Isolation Kit (Human Metabolome Technologies) was used for extracellular vesicle preparation. Briefly, HEK293T cells were transfected with pcDNA3/METTL9-HA, and the culture media were filtered using 0.2 μm membrane filters. Subsequently, extracellular vesicles were isolated using an EVs Quick Filter (#BMA02010) according to the manufacturer’s instructions. The isolated extracellular vesicles (2 μg) were analyzed by Western blotting with anti-HA (Roche, 3F10, 1:1000) and anti-ALIX (abcam, #ab117600, 1:1000) antibodies.

### RT-qPCR

To prepare cDNA templates for real-time RT-PCR, total RNA was extracted from HEK293T, HepG2, HL60, and DMSO-treated HL60 cells using ISOGEN II (Nippon Gene). After DNase treatment, total RNAs were converted to cDNAs using ReverTra Ace qPCR RT Master Mix (TOYOBO) and subjected to real-time quantitative PCR analysis using TB Green Premix EX TaqII (TaKaRa Bio) on a Thermal Cycler Dice (TaKaRa Bio). Relative mRNA expression was calculated using the delta-delta Ct (ΔΔCt) algorithm. The primer set used for qPCR analysis was as follows:

Human *mettl9*_F: 5’-TTGGAGCCAACTAGAGGCAG-3’

Human *mettl9*_R: 5’-CACTTGCCACCTACGTTTTCC-3’

Human β*-actin*_F: 5’-CAAGAGATGGCCACGGCTGC-3’

Human β*-actin*_R: 5’-CTAGAAGCATTTGCGGTGGACG-3’

### Concentration of culture medium

HL60 cells were suspended in serum-free RPMI1640 medium in the presence or absence of PMA (100 nM) and incubated for 1, 2, or 4 h at 37°C in a humidified atmosphere with 5% CO_2_. Then, 500 μL of culture media at each time point was transferred into Amicon Ultra-0.5 mL centrifugal filters (10 K, Merck #UFC501024) and centrifuged at 19,000 × *g* for 30 min. After recovering the concentrated samples, this operation was repeated twice to concentrate 500 μL to 20 μL of the sample. The samples were mixed with 4 μL of 6 × sample buffer solution with reducing reagent (Nacalai tesque #09499-14) and analyzed by Western blotting.

### Plasma collection

Blood samples were collected from the inferior vena cava under deep inhalation anesthesia (isoflurane) using a 23G needle and 1 mL syringe and transferred to tubes containing 10 units of heparin sodium injection (MOCHIDA) as an anticoagulant. Plasma was obtained through two consecutive centrifugation steps. First, the blood was centrifuged at 5,000 rpm for 15 min, and the supernatant was transferred to a new tube and centrifuged again at 5,000 rpm for 15 min to remove platelets and cell debris. The clear supernatants were aliquoted and stored at -80°C.

### Plasma purification

To remove three interfering high-abundance proteins (albumin, IgG, and transferrin) from mouse plasma samples, Multiple Affinity Removal Spin Cartridge Mouse 3 (Agilent #5188-5289) was used according to the manufacturer’s instructions. Briefly, crude mouse plasma (25 μL) from C57BL/6J (4-7 months old) was diluted to 200 μL with Buffer A, followed by filtration through a 0.22 μL spin filter. The diluted plasma sample was added to the spin cartridge and centrifuged at 100 × *g* for 2 min. Furthermore, Buffer A (400 μL) was added to the spin cartridge and centrifuged at 100 × *g* for 3 min. The first and second flow-throw fractions were combined as column-purified plasma, while bound proteins were eluted from the spin cartridge by adding Buffer B (2 mL). This spin cartridge was repeatable after re-equilibration with Buffer A (4 mL).

### *In vitro* methylation assays for mass spectrometry analyses

*In vitro* methylation reactions against plasma proteins were performed using glutathione-sepharose-bound GST-METTL9 WT or 2GA mutant (3 μg) together with column-purified plasma samples in the presence of 10 μM *S*-adenosyl-L-methionine (Sigma-Aldrich #A7007) in PBS. After incubation at 30°C for 6 h, the reactions were concentrated by centrifugation and subjected to LC-MS/MS and MALDI-TOF/MS analyses.

*In vitro* methylation reactions against recombinant proteins were performed with glutathione-sepharose-bound GST-METTL9 WT or 2GA mutant (3 μg) together with GST-cleaved TTR proteins (3 μg), MBP-fused CP fragments (3 μg), or immunoprecipitated full-length CP in the presence of 10 μM *S*-adenosyl-L-methionine (Sigma-Aldrich #A7007) in PBS. After incubation at 30°C for 6 h, the reactions were separated by 14% SDS-PAGE, followed by negative staining with EzStainReverse (ATTO #AE-1310). The band corresponding to TTR was excised, destained with 25 mM Tris-glycine (pH 8.0), and the gel slice was subjected to LC-MS/MS and MALDI-TOF/MS analyses.

### Liquid chromatography-mass spectrometry analysis

After *in vitro* methylation assays, the plasma proteins were directly acid hydrolyzed with 6 M HCl at 110 °C for 24 h, while recombinant proteins in the gel slice were first crushed and passively eluted from the gel slurry using 25 mM Tris-glycine (pH 8.0). After filtration of the gel pieces using ATTO Prep MF cartridges (ATTO #3521370), recombinant proteins were purified from the eluate by methanol-chloroform extraction and subsequently acid hydrolyzed with 6 M HCl at 110 °C for 24 h. Both plasma hydrolysates and recombinant proteins were concentrated by centrifugation, dissolved in 10 μL of water, and injected into a Shimadzu Nexera X2 ultra-high-pressure liquid chromatography-LCMS-8050TM triple-quadrupole mass spectrometer. SeQuant ZIC-HILIC column (2.1 × 150 mm, 3.5 μm; Merck KGaA) with a SeQuant ZIC-HILIC Guard Fitting (1.0 × 14 mm; Merck KGaA) was applied for chromatographic separation. The chromatographic conditions have been described previously [55]. As an internal standard, 25 ng of *N*-propyl-L-arginine (N-PLA) was added to the samples.

MS/MS analysis was performed using collision-induced dissociation (CID) on precursor ions, and the resulting product ions were selectively detected (multiple-reaction monitoring: MRM). Ion intensities were plotted over time, and mass chromatograms were obtained. The optimized instrumental parameters have been described previously [55]. The transitions based on the MRMs of histidine, *N*π-methylhistidine, and *N*τ-methylhistidine were determined to be m/z 156.1 > 110.1, 170.1 > 96.1, and 170.1 > 124.1, respectively. Chromatographic and mass spectrometric system operation, data acquisition, and processing were performed using LabSolutions for LC–MS, version 5.60 software (Shimadzu). Standard amino acid derivatives and N-PLA were diluted in a series ranging from 0.1 to 10 pmol. A calibration curve was generated by plotting the peak area ratios of the analyte to the internal standards against the concentration using linear regression. The concentrations and percentages of analytes observed in each experiment were calculated from the calibration curves.

### Chemical cross-linking assay

HEK293T cells were transfected with pcDNA3/TTR-FLAG WT and point mutants and incubated for 36 h thereafter. For cross-linking intracellular TTR-FLAG, cells were washed twice with PBS and treated with 1 mM disuccinimidyl suberate (DSS; ThermoFisher #21655) diluted in PBS at 4°C for 30 min. After washing with PBS twice, whole-cell lysates were prepared as described above and used for Western blot analysis. For cross-linking secreted TTR-FLAG, the culture media were collected, and cell debris were removed by centrifugation (800 × g for 1 min at 4°C). BS^3^ (bis[sulfosuccinimidyl] suberate, ThermoFisher #21580) was added to the culture medium (final concentration: 1 mM) and incubated for 30 min at room temperature. After quenching with the addition of Tris-HCl (pH 7.6) for 15 min at room temperature (final concentration: 10 mM), samples were mixed with an equal volume of 2 × SDS sample buffer and analyzed by Western blotting.

### Identification of plasma proteins by MALDI-TOF MS

Column-purified mouse plasma proteins were separated by 7%, 11%, and 14% SDS-PAGE, followed by negative staining. The bands corresponding to the ^3^H-labeled bands at approximately 17, 50, and 130 kDa were excised, destained, and subjected to in-gel digestion with trypsin at 37 °C for 24 h after reductive alkylation with DTT and iodoacetamide. Tryptic peptides were eluted from the polyacrylamide gel by sonication in acetonitrile containing 0.1% formic acid. The peptides were desalted with ZipTip μC_18_ (Millipore), and a saturated solution (10 mg/mL) of *R*-cyano-4-hydroxycinnamic acid (Shimadzu GLC) in 0.1% trifluoroacetic acid and 50% acetonitrile was used as the MALDI matrix. MS and MS/MS analyses were performed using an ultrafleXtreme NTA (Bruker). The mass spectrometer was operated in the positive-ion mode and reflector mode under high-voltage conditions as follows: Ion Source1: 20.00 kV, Ion Source2: 17.85 kV, Lens: 8.30 kV, Reflector: 20.80 kV, Reflector2: 11.05 kV. The MS/MS spectra were acquired using the LIFT method under high-voltage conditions as follows: Ion Source1:7.50 kV, Ion Source2: 6.80 kV, Lens: 3.50 kV, Reflector: 29.50 kV, Reflector2: 14.30 kV, LIFT1:19.00 kV, LIFT2: 3.00 kV. The acquired data were processed using FlexAnalysis Version 3.3 (Bruker) and BioTools (Bruker) software. Mascot (Matrix Science) was used as an auxiliary tool to identify plasma proteins, and Swiss-Prot was used as the reference database for protein identification.

### Identification of histidine methylation sites of TTR and CP by MALDI-TOF MS

The gel slice containing TTR and CP proteins was subjected to in-gel digestion with trypsin at 37 °C for 24 h after reductive alkylation with DTT and iodoacetamide. Tryptic peptides were analyzed using MALDI-TOF MS, as described above. The analysis of methylated residues was performed using Mascot and Swiss-Prot.

### Immunofluorescence

HeLa cells were seeded overnight in 35-mm glass-bottom dishes (Matsunami) and transfected with pcDNA3/METTL9-FLAG WT or 2NA mutant for 24 h after seeding. The cells were fixed with 3.7% formaldehyde for 15 min at RT. After washing with PBS three times, the cells were permeabilized with 0.1% Triton X-100 in PBS for 10 min at room temperature, followed by blocking with an immunofluorescence blocking buffer (1 × PBS with 3% BSA and 0.1% Tween20) for 1 h. Each sample was incubated with anti-HA (12CA5, 1:200) and anti-calnexin (Cell Signaling Technology #2679, 1:200) antibodies in a dilution buffer (1 × PBS with 1% BSA and 0.1% Tween20) for 2 h at 4°C. The cells were washed thoroughly with PBS-T (1 × PBS with 0.1% Tween20) and then incubated for 1 h at room temperature with Alexa488 anti-mouse and Alexa594 anti-rabbit antibodies (1:1000) in dilution buffer. After washing with PBS-T, fluorescent signals were acquired using an Olympus FV10i confocal microscope and processed using FLUOVIEW software.

#### In vitro Zinc-binding assay

Zinc-chelating chromatography was performed as previously described [18]. Briefly, an iminodiacetic acid (IDA)-Sepharose column (HiTrap IMAC HP, Cytiva) was equilibrated with zinc sulfate (ZnSO_4_) at a final concentration of 100 mM. After several washes, the TTR peptides (residues 80-114) with either non-methyl- or *N*π-methyl-histidine at His-90 were loaded onto the columns at a concentration of 100 µM, and binding was allowed to proceed for 5 min at room temperature, followed by washing out of the unbound peptides. After extensive washing, the bound peptides were eluted with 50 mM EDTA. The peptide concentrations of both eluted fractions were measured using LC-MS/MS, as described above. The same experiments were independently repeated thrice.

#### Statistical analysis

All studies were performed on at least three independent occasions. The results are plotted as individual values with means. Statistical differences were determined using an unpaired two-tailed t-test for two-group comparisons using GraphPad Prism 9.0 (GraphPad Software). In all figures, statistical significance is indicated by asterisks as follows: *p* < 0.05 (*), *p* < 0.01 (**), and *p* < 0.001 (***).

## Acknowledgements

We thank the members of Fukamizu Laboratory for their helpful discussions. We also thank Rie Sato for her assistance with mass spectrometry analysis and Kae Kumagai for preparing the experimental reagents. HEK293T and HL60 cell lines were kindly provided by the RIKEN BRC and JCRB Cell Bank, respectively. This work was supported by the Japan Society for the Promotion of Science (JSPS) KAKENHI Grant-in-Aid for Scientific Research (B) Grant Numbers JP20H02947 (to H.D.), JP24K01715 (to H.D.), KAKENHI Grant-in-Aid for Scientific Research (A) JP23H00321 (to A.F.), a grant from AMED-CREST Grant Number JP21gm1410010 (to A.F.), and JST SPRING, Japan Grant Number JPMJSP2124 (to N.I.).

## Author contributions

Conceptualization, H.D. and A.F.; methodology, H.D., N.I., N.O., N.S., R.T., R.K., S.H., K.M., and K.K.; Investigation, H.D., N.I., N.O., N.S., R.T., R.K., S.H., and K.K.; writing—original draft, H.D.; writing—review & editing, N.I. and A.F.; funding acquisition, H.D., N.I., and A.F.; resources, K.M.; supervision, H.D. and A.F.

