## Supplementary Figures for "Secreted protein methyltransferase METTL9 catalyzes *N*π-histidine methylation of extracellular plasma proteins"

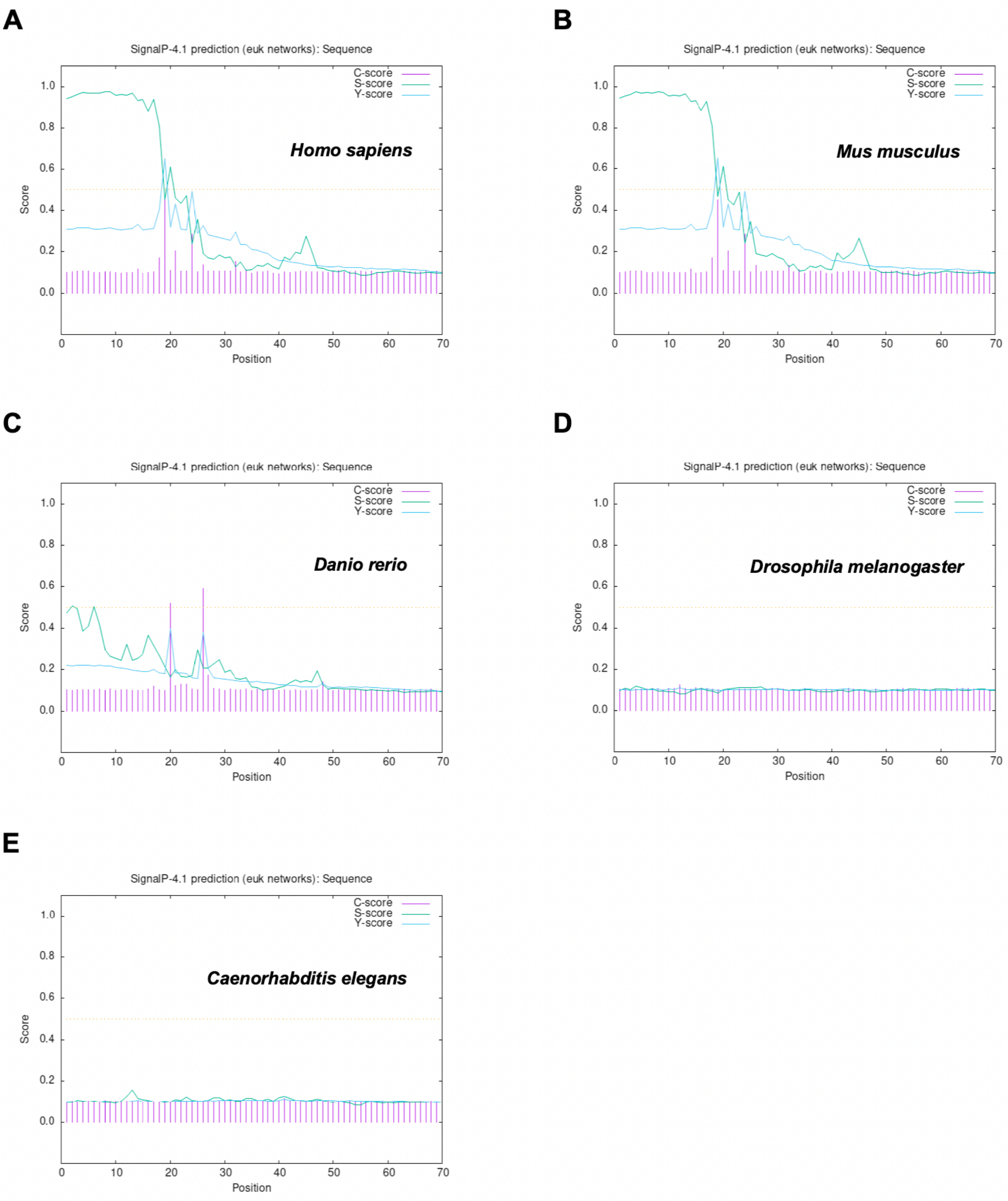


**Figure S1. Prediction of signal peptide sequences of METTL9 orthologs**

(**A**-**E**) SignalP 4.1 server predicted the presence and location of signal peptide cleavage sites in amino acid sequences from METTL9 orthologs. Sequence alignment of the N-terminal end of METTL9 orthologs across species. *Homo sapiens* (**A**), *Mus musculus* (**B**), *Danio rerio* (**C**), *Drosophila melanogaster* (**D**), *and Caenorhabditis elegans* (**E**). C-score: raw cleavage site score; S-score: signal peptide score; Y-score: combined cleavage site score.

**A**

**Protein View: CALX_HUMAN**

**Protein sequence coverage: 42%**

1 MEGKWLLCML LVLGTAIVEA HDGHDDDVID IEDDLDDVIE EVEDSKPDTT APPSSPKVTY K**APVPTGEVY FADSFDRGTL**

81 **SGWILSK**AKK **DDTDDEIAKY DGKWEVEEMK** ESK**LPGDKGL VLMSRAKHHA ISAK**LNKPFL FDTKPLIVQY EVNFQNGIEC

161 GGAYVK**LLSK TPELNLDQFH DKTPYTIMFG PDK**CGEDYK**L HFIFR**HKNPK **TGIYEEKHAK RPDADLKTYF TDKK**THLYTL

241 ILNPDNSFEI LVDQSVVNSG NLLNDMTPPV NPSR**EIEDPE DRKPEDWDER PKIPDPEAVK PDDWDEDAPA K**IPDEEATKP

321 EGWLDDEPEY VPDPDAEKPE DWDEDMDGEW EAPQIANPRC ESAPGCGVWQ RPVIDNPNYK GK**WKPPMIDN PSYQGIWKPR**

401 **KIPNPDFFED LEPFR**MTPFS AIGLELWSMT SDIFFDNFII CADR**RIVDDW ANDGWGLKK**A ADGAAEPGVV GQMIEAAEER

481 PWLWVVYILT VALPVFLVIL FCCSGK**KQTS GMEYKK**TDAP QPDVK**EEEEE KEEEK**DKGDE EEEGEEK**LEE KQK**SDAEEDG

561 GTVSQEEEDR KPK**AEEDEIL NRSPR**NRKPR RE

**B**

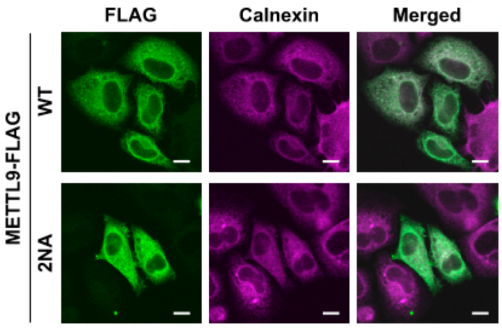


**Figure S2. Calnexin is identified as an METTL9-binding protein**

(**A**) MASCOT search results for METTL9-binding proteins. The matched peptides are shown in bold red. (**B**) Intracellular localization of METTL9 WT and 2NA mutant. Representative immunofluorescence images of METTL9 WT (top) or 2NA mutant (bottom) localization in HeLa cells are shown. Green, METTL9-FLAG; magenta, Calnexin. Images of METTL9-FLAG were merged with calnexin staining and shown as white pixels. The scale bars represent 10 μM.

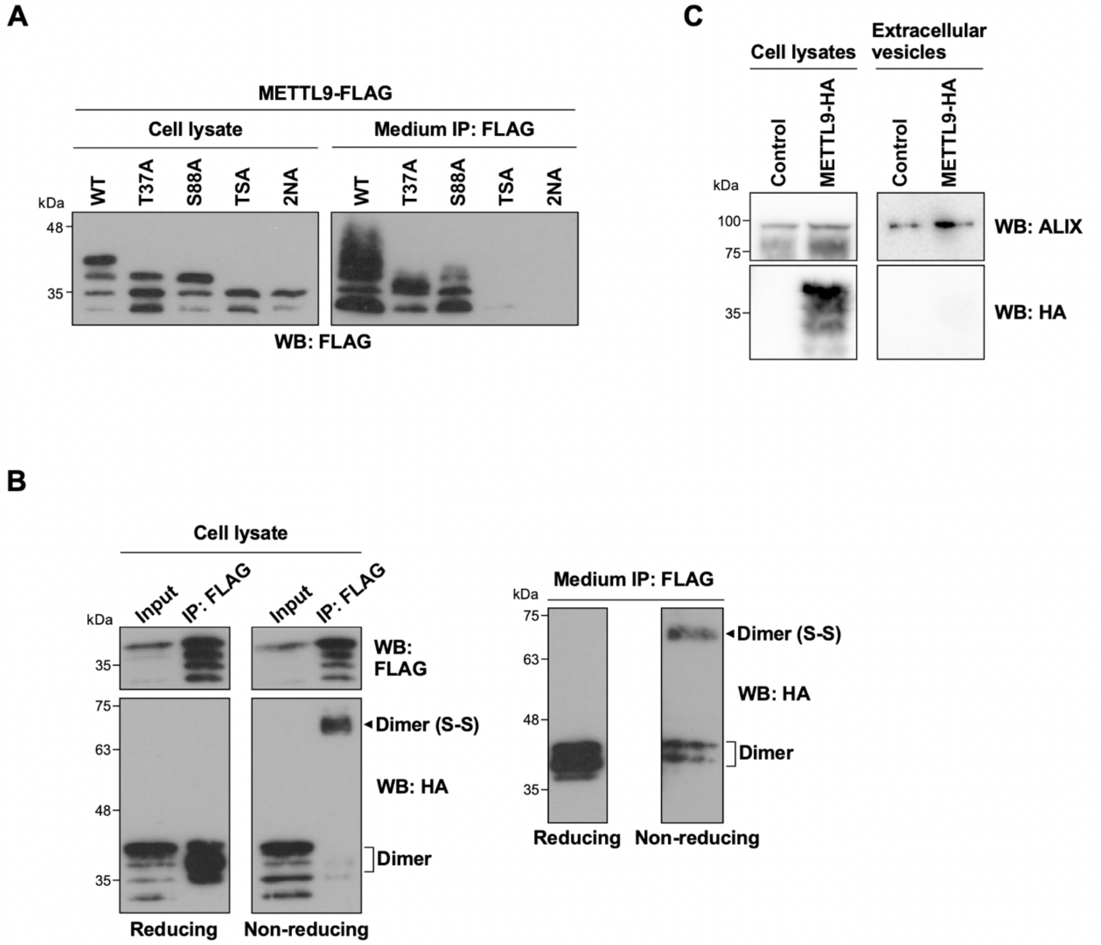


**Figure S3. N-glycosylated METTL9 is secreted extracellularly as a disulfide dimer**

(**A**) Medium immunoprecipitation of METTL9. The culture media of cells expressing METTL9-FLAG WT and several mutants, as indicated, were immunoprecipitated with anti-FLAG antibody. These immunoprecipitates and cell lysates were analyzed by Western blotting with anti-FLAG antibody. (**B**) Extracellular dimerization of METTL9 via disulfide bond formation. Cell lysates (*Left*) and culture media (*Right*) of cells co-expressing FLAG- and HA-tagged METTL9 were immunoprecipitated with an anti-FLAG antibody under reducing or non-reducing conditions. The immunoprecipitates were analyzed by Western blotting with anti-FLAG and anti-HA antibodies. The arrowhead indicates a band of METTL9 dimers formed via disulfide bond formation. The input indicates 2% of the whole-cell lysates used for immunoprecipitation. (**C**) Extracellular vesicles do not contain METTL9. The culture media of cells expressing METTL9-HA were purified using the EVs Quick Filter. The isolated exosome fraction (*Right*) and cell lysates (*Left*) were analyzed by Western blotting with anti-ALIX and anti-HA antibodies.

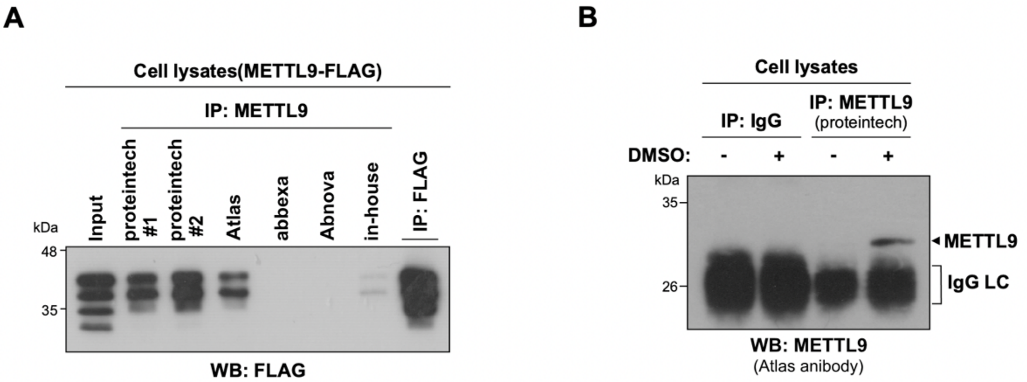


**Figure S4. Immunoprecipitation of overexpressed and endogenous METTL9**

(**A**) Verification of the immunoprecipitation efficiency of anti-METTL9 antibodies. Cell lysates expressing METTL9-FLAG were immunoprecipitated with commercially available or in-house anti-METTL9 antibodies, as indicated, or with anti-FLAG antibodies. These immunoprecipitates were analyzed by Western blotting with an anti-FLAG antibody. The input indicates 2% of the whole-cell lysates used for immunoprecipitation. (**B**) Endogenous METTL9 expression in HL60 cells. Lysates of HL60 cells treated with or without DMSO were immunoprecipitated with normal rabbit IgG or anti-METTL9 antibody (proteintech) and then analyzed by Western blotting with anti-METTL9 antibody (Atlas). An arrowhead indicates a band of endogenous METTL9.

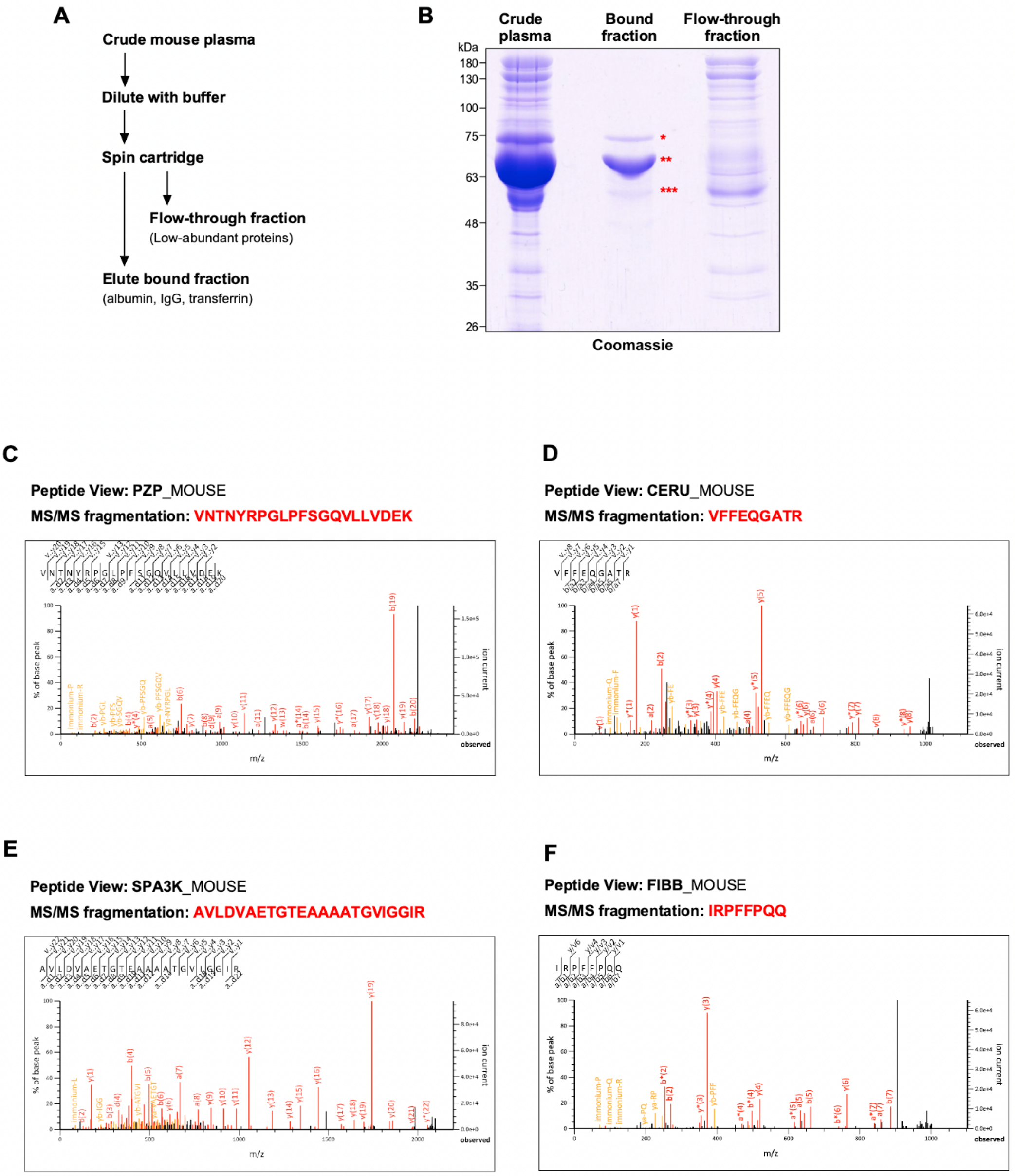
A
Crude mouse plasma
Dilute with buffer
Spin cartridge
Flow-through fraction
(Low-abundant proteins)
Elute bound fraction
(albumin, IgG, transferrin)
B
Crude plasma
Bound fraction
Flow-through fraction
kDa
180
130
100
75
63
48
35
26
*
**
***
Coomassie
C
Peptide View: PZP_MOUSE
MS/MS fragmentation: VNTNYRPGLPFSGQVLLVDEK
D
Peptide View: CERU_MOUSE
MS/MS fragmentation: VFFEQGATR
E
Peptide View: SPA3K_MOUSE
MS/MS fragmentation: AVLDVAETGTEAAAATGVIGGIR
F
Peptide View: FIBB_MOUSE
MS/MS fragmentation: IRPFFPQQ
% of base peak
m/z
Ion current

**G**

**Peptide View:** VTDB_MOUSE
**MS/MS fragmentation:** RTQVPEVFLSK

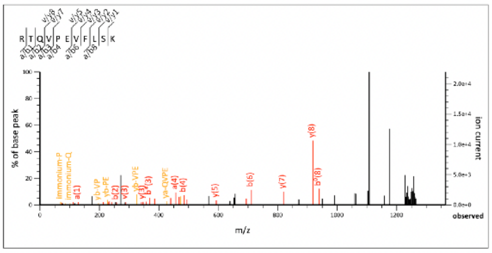


**H**

**Peptide View:** A1AT1_MOUSE
**MS/MS fragmentation:** TLMSPLGITR

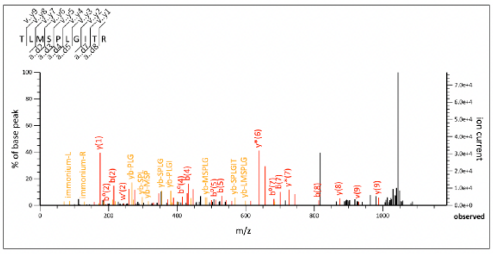


**I**

**Peptide View:** FIBG_MOUSE
**MS/MS fragmentation:** ESGLYFIRPLK

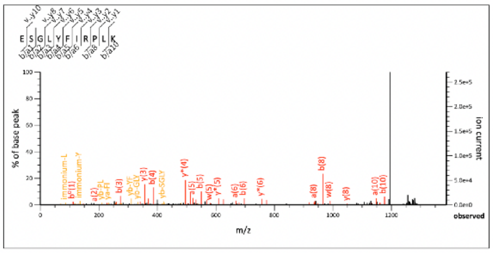


**J**

**Peptide View:** TTHY_MOUSE
**MS/MS fragmentation:** VLDAVRGSPAVDVAVK

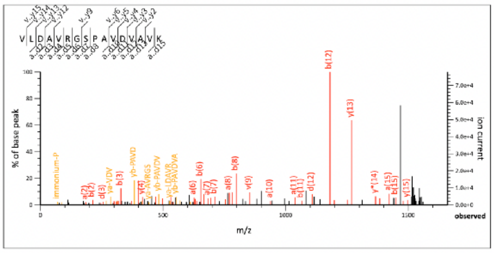


**K**

**Ceruloplasmin**_MOUSE

1 MKFLLLSTFI FLYSSLALAR DKHYFIGITE AVWDYASGTE EKKLISVDTE QSNFYLQNGP DRIGRKYKKA LYFEYTDGTF
81 SKTIDKPAWL GFLGPVIKAE VEDKVYVHLK NLASRIYTF**H** **AH**GVTYTKEY EGAVYPDNTT DFQRADDKVL PGQQYVYVLH
161 ANEPSPGEGD SNCVTRIY**HS** **H**VDAPKDIAS GLIGPLILCK KGSLYKEKEK NIDQEFVLMF SVVDENLSWY LEDNIKTFCS
241 EPEKVDKDNE DFQESNRMYS INGYTFGSLP GLSMCAADRV KWYLFGMGNE VDVHSAFFHG QALTSRNYQT DIINLFPATL
321 IDAYMVAQNP GVWMLSCQNL NHLKAGLQAF FQVRDCNKPS PEDNIQDRHV RHYYIAAEEV IWNYAPSGTD IFTGENLTAL
401 ESDSRVFFEQ GATRIGGSYK KMAYREYTDG SFTNRKQRGP DEEHLGILGP VIWAEVGDTI KVTFHNKGQH PLSIQPMGVS
481 FTAENEGTYY GPPGRSSQQA ASHVAPKETF TYEWTVPKEM GPTYADPVCL SKMYYSGVDP TKDIFTGLIG PMKICKKGSL
561 LADGRQKDVD KEFYLFPTVF DENESLLLDD NIRMFTTAPD QVDKEDEDFQ ESNKMHSMNG FMYGNQPGLN MCLGESIVWY
641 LFSAGNEADV HGIYFSGNTY LSKGERRDTA NLFPHKSLTL LMNPDTKGTF DVECLTTDHY TGGMKQKYTV NQCQRQFEDF
721 TVYLGERTYY VAAVEVEWDY SPSRAWEKEL HHLQEQNVSN VFLDKEEFFI GSKYKKVVYR QFTDSSFREQ VKRRAEDEHL
801 GILGPPIHAN VGDKVKVVFK NMATRPYSI**H** **AH**GVKTESST VVPTLPGEVR TYTWQIPERS GAGREDSACI PWAYYSTVDR
881 VKDLYSGLIG PLIVCRKSYV KVFSPKKKME FFLLFLVFDE NESWYLDDNI KTYSEHPEKV NKDNEEFLES NKMHAINGKM
961 FGNLQGLTMH VKDEVNWYVM GMGNEIDLHT V**HFHGH**SFQY KHRGVYSSDV FDLFPGTYQT LEMFPQTPGT WLL**HCH**VTD**H**
1041 **VH**AGMATTYT VLPVEQETKS G

**L**

**Transthyretin**_MOUSE

1 MASLRLFLLC LAGLVFVSEA GPAGAGESKC PLMVKVLDAV RGSPAVDVAV KVFKKTSEGS WEPFASGKTA ESGELHGLTT
81 DEKFVEGVYR VELDTKSYWK TLGISPFHEF ADVVFTANDS G**HRH**YTIAAL LSPYSYSTTA VVSNPQN

**Figure S5. Affinity removal of high-abundant proteins from mouse plasma**

(**A**) Schematic of the multiple affinity removal system for mouse plasma (according to the instructions of Agilent Technologies). (**B**) Verification of the removal of high-abundance proteins from mouse plasma. Crude mouse plasma, column-bound fractions, and flow-through fractions were detected using Coomassie blue staining after SDS-PAGE. Red asterisks indicate bands corresponding to transferrin (*), albumin (**), and IgG (***). (**C**-**J**) MS/MS fragmentation spectra of mouse PZP (**C**), CERU (**D**), SPA3K (**E**), FIBB (**F**), VTDB (**G**), A1AT1 (**H**), FIBG (**I**), and TTHY (**J**). The matched peptides are shown in bold red. The b-series and y-series peptide fragment ions are shown. (**K** and **L**) Amino acid sequences of mouse ceruloplasmin (**K**) and transthyretin (**L**). The consensus motif for METTL9-dependent methylation (H-X-H) is shown in bold red.

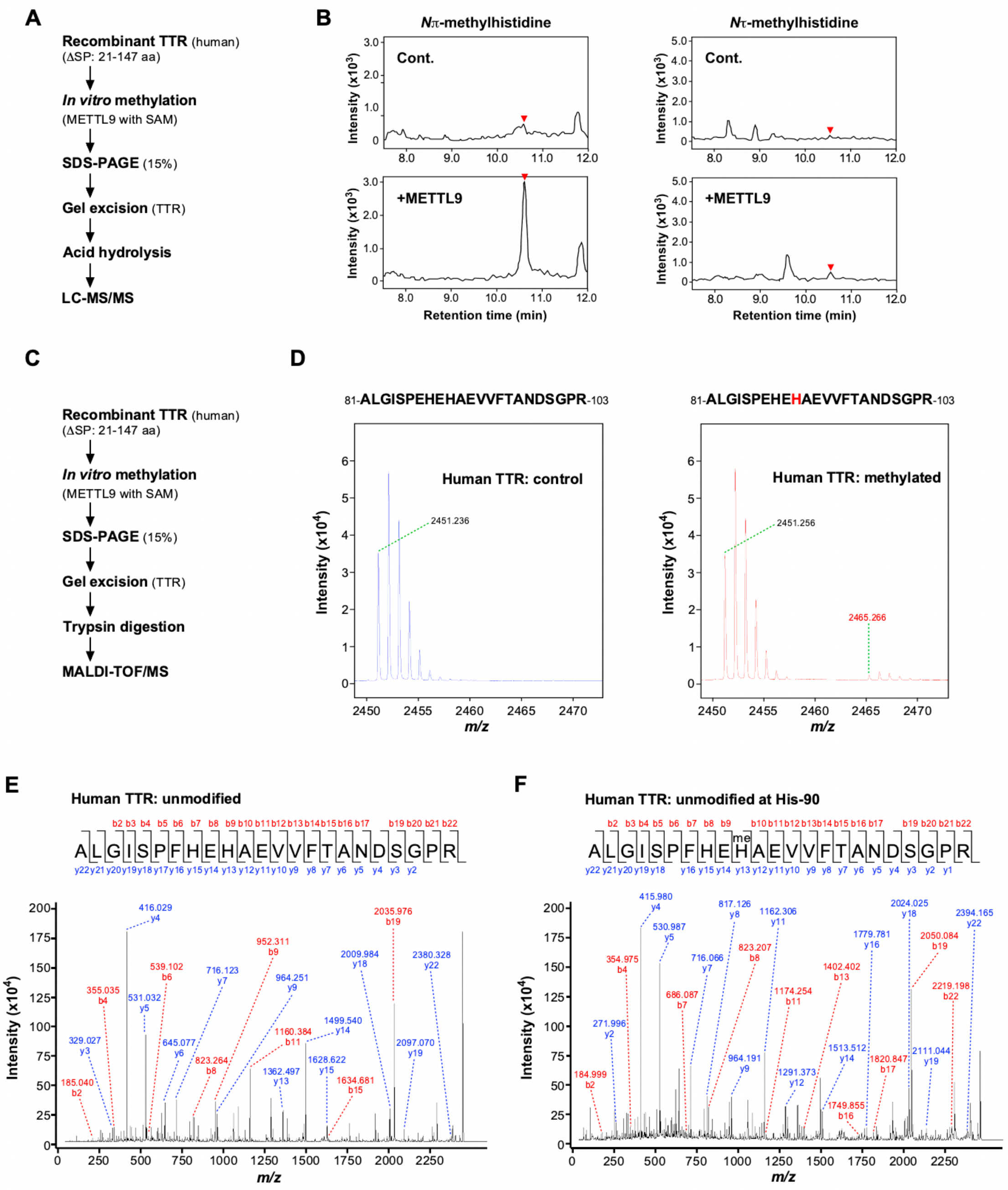


**Figure S6. Identification of METTL9-mediated methylation site in human TTR**

(**A**) Schematic representation of the strategy used to measure the levels of histidine methylation in human TTR after *the in vitro* methylation assay. (**B**) METTL9 catalyzes *N*π-methylhistidine formation in human TTR *in vitro*. Recombinant TTR was incubated with recombinant METTL9 WT in the presence of S-adenosylmethionine (SAM). After SDS-PAGE, the methylhistidine content of TTR was determined by LC-MS/MS and is shown as chromatograms. Red arrowheads indicate the retention times of *N*π- and *N*τ-methylhistidine defined by their standards. (**C**) Schematic representation of the strategy to identify methylation sites of human TTR after *in vitro* methylation assay. (**D**) Tryptic peptides of human TTR contain monomethylated amino acids. *In vitro* methylation reactions were resolved using SDS-PAGE, followed by reverse staining. The portion of the gel corresponding to TTR was digested with trypsin and analyzed using MALDI-TOF/MS. Unmodified (*Left*) and monomethylated (*Right*) peptide sequences from human TTR are shown. The METTL9-mediated methylation site at His-90 is denoted in red. (**E** and **F**) MS/MS fragmentation spectra showing monomethylation of human TTR at His-90. The 23-amino acid peptide sequence and b (red) and y (blue) series peptide fragment ions for the unmodified (**E**) and methylated (**F**) peptide products are shown.

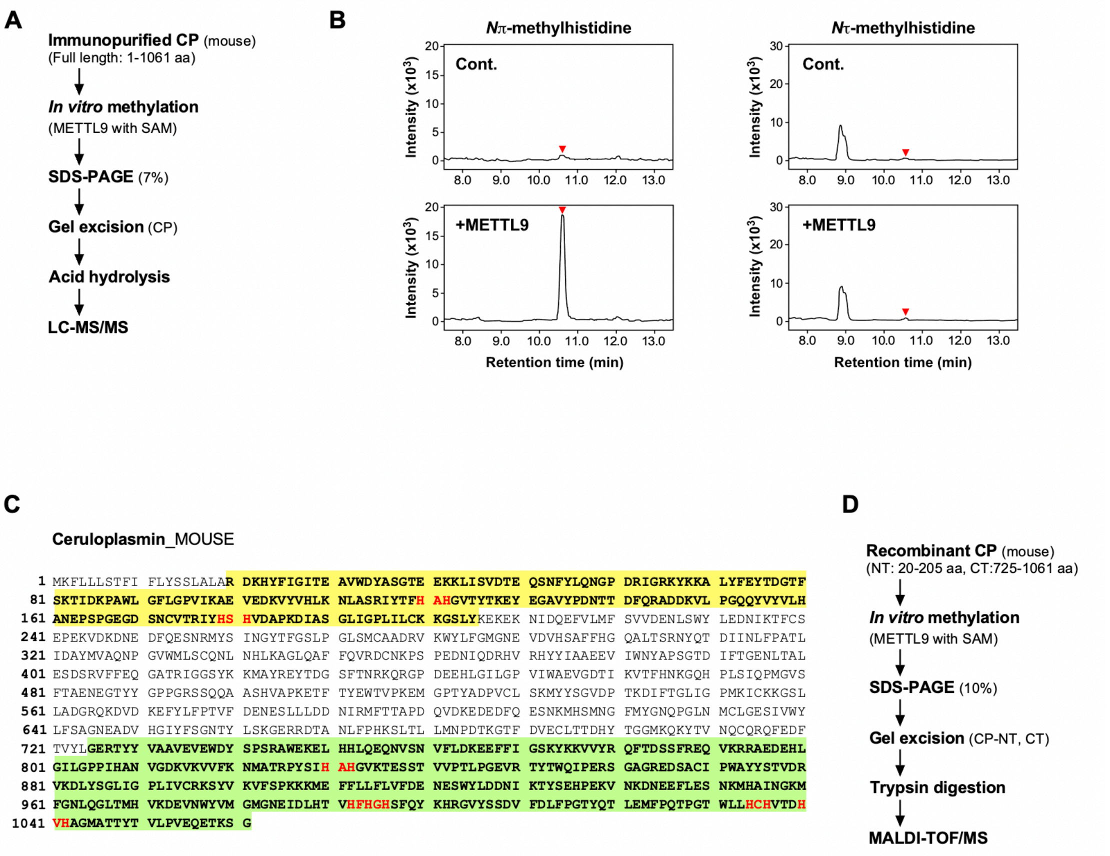
A
Immunopurified CP (mouse)
(Full length: 1-1061 aa)
In vitro methylation
(METTL9 with SAM)
SDS-PAGE (7%)
Gel excision (CP)
Acid hydrolysis
LC-MS/MS
B
Nπ-methylhistidine
Nτ-methylhistidine
Cont.
+METTL9
Intensity (x10^3)
Retention time (min)
C
Ceruloplasmin_MOUSE
1 MKFLLLSTFI FLYSSLALAR DKHYFIGITE AVWDYASGTE EKKLISVDTE QSNFYLQNGP DRIGRKYKKA LYFEYTDGTF
81 SKTIDKPAWL GFLGPVIKAE VEDKVYVHLK NLASRIYTFH AHGVTYTKEY EGAVYPDNTT DFQRADDKVL PGQQYVYVLH
161 ANEPSPGEGD SNCVTRIYHS HVDAPKDIAS GLIGPLILCK KGSLYKEKEK NIDQEFVLMF SVVDENLSWY LEDNIKTFCS
241 EPEKVDKDNE DFQESNRMYS INGYTFGSLP GLSMCAADRV KWYLFGMGNE VDVHSAFFHG QALTSRNYQT DIINLFPATL
321 IDAYMVAQNP GVWMLSCQNL NHLKAGLQAF FQVRDCNKPS PEDNIQDRHV RHYYIAAEEV IWNYAPSGTD IFTGENLTAL
401 ESDSRVFFEQ GATRIGGSYK KMAYREYTDG SFTNRKQRGP DEEHLGILGP VIWAEVGDTI KVTFHNKGQH PLSIQPMGVS
481 FTAENEGTYY GPPGRSSQQA ASHVAPKETF TYEWTVPKEM GPTYADPVCL SKMYYSGVDP TKDIFTGLIG PMKICKKGSL
561 LADGRQKDVD KEFYLFPTVF DENESLLLDD NIRMFTTAPD QVDKEDEDFQ ESNKMHSMNG FMYGNQPGLN MCLGESIVWY
641 LFSAGNEADV HGIYFSGNTY LSKGERRDTA NLFPHKSLTL LMNPDTKGTF DVECLTTDHY TGGMKQKYTV NQCQRQFEDF
721 TVYLGERTYY VAAVEVEWDY SPSRAWEKEL HHLQEQNVSN VFLDKEEFFI GSKYKKVVYR QFTDSSFREQ VKRRAEDEHL
801 GILGPPIHAN VGDKVKVVFK NMATRPYSIH AHGVKTESST VVPTLPGEVR TYTWQIPERS GAGREDSACI PWAYYSTVDR
881 VKDLYSGLIG PLIVCRKSYV KVFSPKKKME FFLLFLVFDE NESWYLDDNI KTYSEHPEKV NKDNEEFLES NKMHAINGKM
961 FGNLQGLTMH VKDEVNWYVM GMGNEIDLHT VHFHGHSFQY KHRGVYSSDV FDLFPGTYQT LEMFPQTPGT WLLHCHVTDH
1041 VHAGMATTYT VLPVEQETKS G
D
Recombinant CP (mouse)
(NT: 20-205 aa, CT:725-1061 aa)
In vitro methylation
(METTL9 with SAM)
SDS-PAGE (10%)
Gel excision (CP-NT, CT)
Trypsin digestion
MALDI-TOF/MS

**E**

**Peptide View: CERU**_MOUSE; **unmodified**

**MS/MS fragmentation: IYTFHAHGVTYTK**

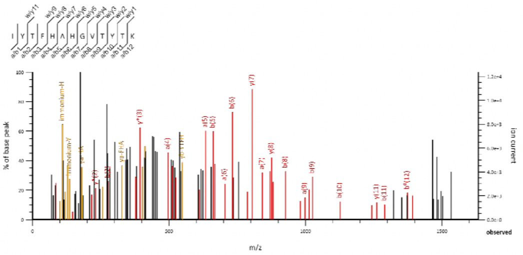

**F**

**Peptide View: CERU**_MOUSE; **methylated at His-122**

**MS/MS fragmentation: IYTFHAHGVTYTK**

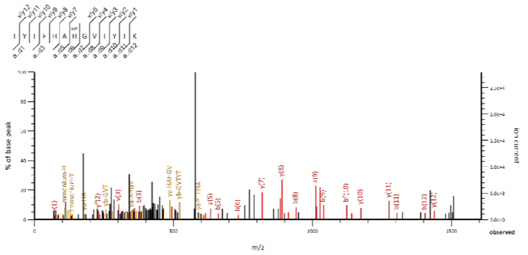

**G**

**Peptide View: CERU**_MOUSE; **unmodified**

**MS/MS fragmentation: IYHSHVDAPK**

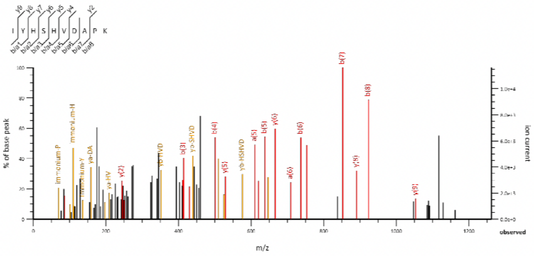

**H**

**Peptide View: CERU**_MOUSE; **methylated at His-181**

**MS/MS fragmentation: IYHSHVDAPK**

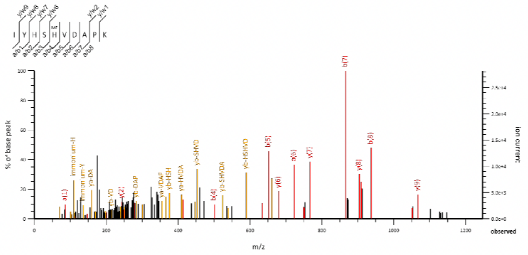

**I**

**Peptide View: CERU**_MOUSE; **unmodified**

**MS/MS fragmentation: NMATRPYSIHAHGVK**

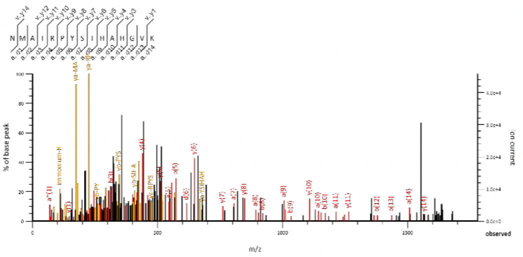

**J**

**Peptide View: CERU**_MOUSE; **methylated at His-832**

**MS/MS fragmentation: NMATRPYSIHAHGVK**

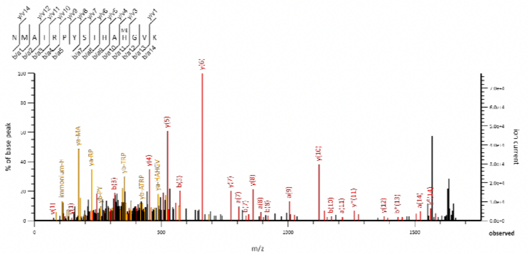

**Figure S7. Identification of METTL9-mediated methylation site in mouse CP**

(**A**) Schematic of the strategy used to measure the levels of histidine methylation in mouse CP after *the in vitro* methylation assay. (**B**) METTL9 catalyzes *N*π-methylhistidine formation in mouse CP *in vitro*. Immunopurified CP was incubated with recombinant METTL9 WT in the presence of SAM. After SDS-PAGE, the methylhistidine content of CP was determined by LC-MS/MS and is shown as chromatograms. Red arrowheads indicate the retention times of *N*π- and *N*τ-methylhistidine defined by their standards. (**C**) Amino acid sequence of the mouse CP. The N-terminal (residues 20-205) and C-terminal (residue 725-1061) regions of CP are denoted in bold and highlighted in yellow and green, respectively. The consensus motif for METTL9-dependent methylation (H-X-H) is shown in bold red. (**D**) Schematic representation of the strategy used to identify methylation sites of mouse CP after *in vitro* methylation assay. (**E** and **F**) MS/MS fragmentation spectra showing monomethylation of mouse CP at His-122. Matched peptides are shown in bold red, and methylated histidine residues are underlined. The b-series and y-series peptide fragment ions for unmodified (**E**) and methylated (**F**) peptide products are shown. (**G** and **H**) MS/MS fragmentation spectra showing monomethylation of mouse CP at His-181. Matched peptides are shown in bold red, and methylated histidine residues are underlined. The b-series and y-series peptide fragment ions for the unmodified (**G**) and methylated (**H**) peptide products are shown. (**I** and **J**) MS/MS fragmentation spectra showing monomethylation of mouse CP at His-832. Matched peptides are shown in bold red, and methylated histidine residues are underlined. The b-series and y-series peptide fragment ions for unmodified (**I**) and methylated (**J**) peptide products are shown.
